# Investigation of allele-specific expression in citrus hybrids reveals the association between siRNA-mediated de novo methylation and high expression in citrus genomes

**DOI:** 10.1101/2025.07.25.666733

**Authors:** Diop Khadidiatou, Bonnin Marie, Gibert Anaïs, Christel Llauro, Yann Froelicher, Hufnagel Barbara, Picault Nathalie, Pontvianne Frédéric

## Abstract

DNA methylation plays a central role in the regulation of gene expression. In plants, methylation occurs in the CG, CHG and CHH contexts, *via* distinct DNA methyltransferases including MET1, CMT3 and the RNA-directed DNA Methylation (RdDM) pathway *via* DRM2. In interspecific hybrids, these epigenetic mechanisms are confronted to a mixed small RNA population and two subgenomes harbouring specific methylation patterns, therefore generating unique expression profiles. The aim of this work was to understand these regulations by analysing gene expression, DNA methylation and small RNAs in a Citrus hybrid resulting from the cross between *C. reticulata* (mandarin) and *C. australasica* (finger lime). Haplotype-resolved subgenomes assembly identified hundreds of allele-specifically expressed genes. Asymmetric reprogramming of methylation was observed, in particular an increase in CHH in *C. australasica* haplotype. Surprisingly, CHH methylation, usually associated with gene silencing, was correlated here with increased expression, but also 24nt small RNA populations at their promoter regions. Similar analyses of the parental lines and other citrus species suggest the correlation between CHH methylation-enriched promoter and high expression level is not due to the hybridization, but seem to be generally true for all citrus. These observations suggest that, in citrus fruit, RdDM could activate transcription. This work also provides a full pipeline to analyse the expression profiles and DNA methylation in complex hybrids, which could be crucial for anticipating varieties resistant to diseases and the current threats affecting citriculture such as the Huanglongbing disease.

## INTRODUCTION

Citrus fruits (*Citrus spp*.) are among the most economically and culturally significant fruit crops worldwide, with global production estimated at approximately 140–146 million tons annually (Mukhametzyanov et al., 2024). Their exceptional phenotypic diversity in fruit size, colour, flavour, essential oil composition, and nutritional value originates from a limited number of sexually compatible ancestral taxa - *C. medica*, *C. maxima*, *C. reticulata*, *C. micrantha*, and *C. cavaleriei* - whose interspecific hybridizations, followed by clonal propagation *via* apomixis and grafting, gave rise to the multitude of cultivated varieties ((Wu et al. 2018);(Ollitrault et al. 2020);(Curk et al. 2022)). While most commercial cultivars derive from combinations of these Asian ancestral taxa, recent phylogenomic analyses have led to the integration of formerly distinct genera such as *Microcitrus* and *Eremocitrus* into the genus *Citrus* ((Wu et al. 2018);(Curk et al. 2022)). These Oceanian lineages, including *C. australasica*, are genetically more distant from the Asian citrus gene pool, yet remain largely unused in modern breeding programs despite their potential to introduce novel adaptive traits (Nakandala et al. 2023).

Modern citriculture now faces an alarming convergence of challenges that threaten both productivity and sustainability. Among these, Huanglongbing (HLB), caused by *Candidatus Liberibacter spp*. and vectored by psyllids, has devastated citrus production in major growing regions, including Brazil, the United States, and China ((Costa et al. 2021);(Alquézar et al. 2022)). Alongside HLB, producers must cope with the intensification of abiotic stresses such as drought, salinity, and thermal extremes, as well as an increasing burden of viral and fungal pathogens ((Yesiloglu et al. 2022);(Dahro et al. 2023)). These pressures are exacerbated by the narrow genetic base of most cultivated varieties, which limits the availability of resilient alleles within the traditional gene pool. As a result, breeding efforts are increasingly turning toward exotic and underutilized germplasm to overcome these constraints and ensure long-term sustainability of citrus production ((Al-Khayri 2018);(Cimen et al. 2022)).

In this context, wide interspecific hybridization within the expanded *Citrus* genus offers a promising avenue for enhancing the adaptive potential of cultivated citrus. Crosses involving phylogenetically distant species such as *C. australasica* (formerly *Microcitrus*), *C. glauca* (formerly *Eremocitrus*), or *C. trifoliata* (formerly *Poncirus*) now all taxonomically integrated within *Citrus* have yielded germplasm with unique traits, including drought tolerance, cold hardiness, and partial HLB tolerance ((Peng et al. 2020);(Alves et al. 2022);(Mahmoud et Dutt 2025)). Despite their potential, these lineages remain largely untapped in commercial breeding programs. Notably, *C. australasica* diverged from the main Asian citrus lineages during a distinct radiation event in the early Pliocene, approximately 4 million years ago (Wu et al. 2018), making it a particularly informative model to investigate the genomic consequences of wide hybridization, which integrates both authentic post-hybridization regulatory changes and the persistence of conserved intra-genus patterns.

Chromatin modifications, and especially DNA methylation, can impact gene expression *via* different pathways. In plants, DNA methylation occurs in three cytosine sequence contexts: CG, CHG, and CHH (where H = A, T, or C). CG methylation is maintained primarily by METHYLTRANSFERASE 1 (MET1), while CHG methylation depends on CHROMOMETHYLASE 3 (CMT3). CHH methylation is guided by both the RNA-directed DNA methylation (RdDM) pathway, involving DOMAINS REARRANGED METHYLTRANSFERASE 2 (DRM2), and the CMT2 pathway. These distinct but partially overlapping mechanisms ensure the establishment and maintenance of methylation across the genome, usually with a repressive role although it can vary depending on the context (Law et Jacobsen 2010). Together with other chromatin modifications such as histone post-translational modifications, DNA methylation patterns shape the epigenome. In the case of hybrids, it is not just two different genomes that are associated, but two different epigenomes, thus carrying both genetic and epigenetic variation.

Interspecific hybridization, is a common process in plants which enables the apparition of new phenotypes and species (Mason et Batley 2015). In breeding, this technique is widely used especially when hybrid vigor, also known as heterosis, is displayed in the F1 sibling. Heterosis is usually characterized in F1 by an increased biomass production and stress tolerance (Birchler et al. 2010). Recent sequencing approaches revealed that hybridization could be considered as a genomic and an epigenomic shock that, *in fine*, lead to the establishment of an hybrid specific transcriptional program which is not simply an addition of each parental transcriptional program (Hochholdinger et Yu 2025). In *Arabidopsis thaliana*, in rice, in maize and in cotton, DNA methylation and siRNA populations are altered in the hybrid. Changes to the hybrid methylome can occur in a way that differential methylation patterns inherited from the parent could be compensated to resemble each other, potentially *via* the action of small RNA (Greaves et al. 2015). However, this observation needs more evidence.

Creating hybrid resistant to pathogens, with a fruit pomology reaching the standard for citriculture is necessary to save a sector in great danger due to the various threats, in particular to diseases such the aforementioned HLB. The aim of the present study is therefore not to identify resistance mechanisms per se, but to establish a haplotype-resolved framework to characterize allele-specific gene expression, DNA methylation, and small RNA accumulation in a complex interspecific citrus hybrid. To better understand these regulatory pathways, we analysed the expression profiles in a hybrid between two species separated by more than 4 million years of evolution, *Citrus reticulata* (mandarin) and *Citrus* (formerly *Microcitrus*) *australasica* (finger lime). In addition, to determine the impact of modifications to epigenetic signatures such as DNA methylation on this expression profile, we analysed the overall methylation of the genomes, but also of the associated small RNAs that enable them to be deposited in certain contexts. In the end, this study shows that these pathways participate in the establishment of differential allelic expression within the hybrid, but also that de novo methylation seems to play a particular role in citrus.

## RESULTS

### Assembly of an inter-specific citrus hybrid with a haplotype resolution

Although no direct genome comparison was performed prior to assembly, the two parental lines (*C. reticulata* and *C. australasica*) are known from phylogenetic studies to have diverged more than 4 million years ago (Wu et al. 2018). This long evolutionary separation suggests substantial sequence divergence between their genomes, likely including structural variation and high heterozygosity in their hybrid offspring. We therefore performed haplotype-resolved assembly rather than a conventional consensus assembly as previously published as trio-binning (Koren et al. 2018). We generated long-read sequencing using Oxford Nanopore Technologies of the hybrid individual with a 160X genome coverage using leaf tissue. To enable the assignment of hybrid long reads to their respective parental origins prior to assembly, we also obtained parental-specific k-mers from Illumina Tell-Seq libraries generated for each parent, as well short-read sequencing. This strategy allowed the realization of accurate trio-binning and independent reconstruction of both haplotypes (Figure 1A). Each assembled haplotypes consisted of only nine contigs, corresponding to the 9 chromosomes with a total assembly size of 349 Mb for *C. reticulata* and 359 Mb for *C. australasica*. The assemblies were highly contiguous (N50 = 38.0 Mb and 39.9 Mb, respectively; L50 = 4 for both) and exhibited consistent GC content (∼34%).

**Figure 1.**
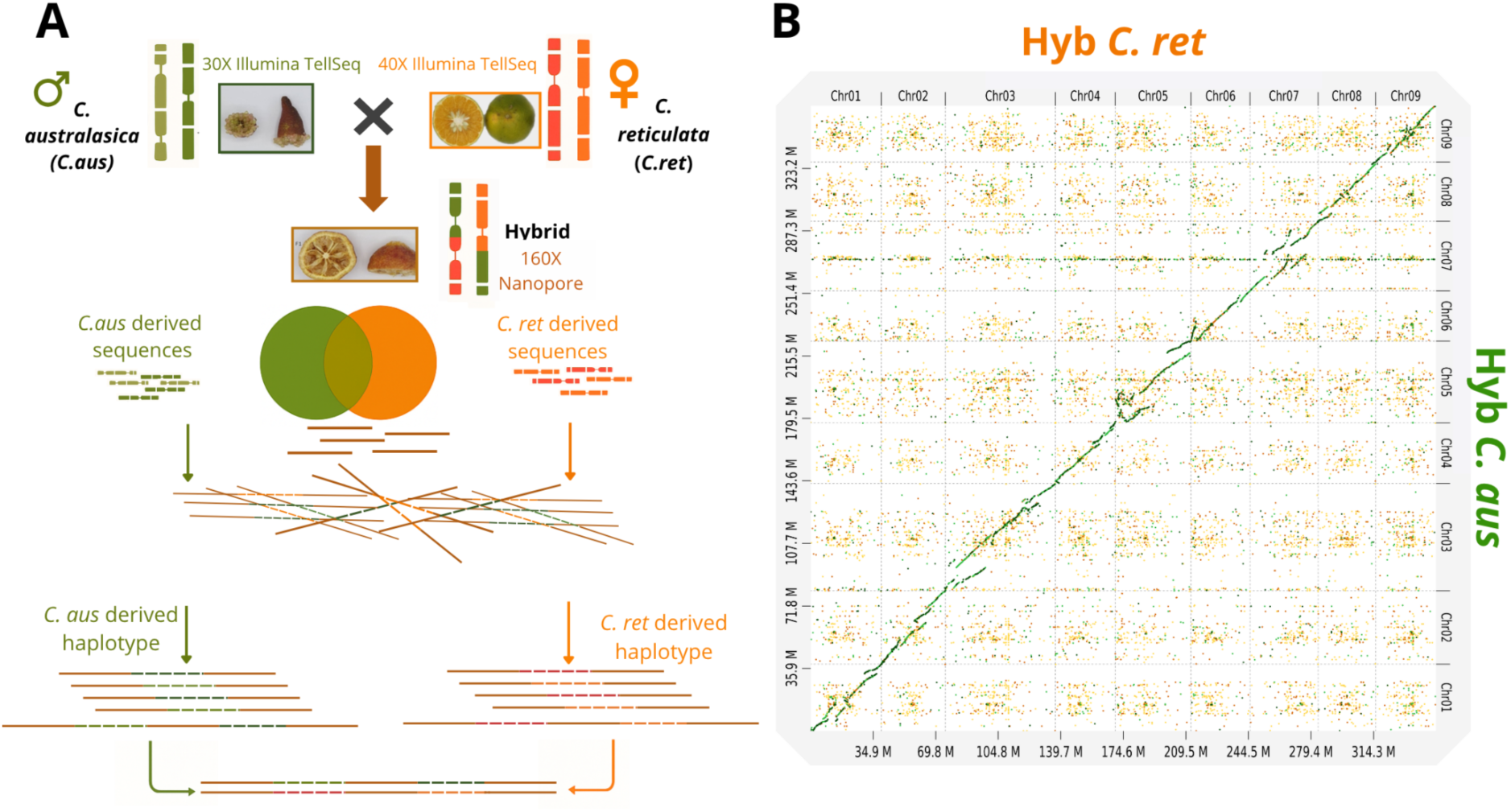
Haplotype-resolved genome assembly and structural variation. (A) Overview of the trio-binning genome assembly strategy used to generate haplotype-resolved assemblies of the citrus hybrid. Long reads from the hybrid were partitioned based on parental short-read data to generate separate assemblies corresponding to the maternal (C. reticulata, orange) and paternal (C. australasica, green) haplotypes. (B) Dot-plot alignment between the maternal (Hyb C. ret) and paternal (Hyb C. aus) genome assemblies showing overall synteny and structural rearrangements. Diagonal lines indicate homologous regions; off-diagonal signals reflect structural variations between haplotypes.

To assess the assembly quality of both haplotypes, we anchored their pseudomolecules to a consensus *Citrus* genetic map (Ollitrault et al. 2024), constructed from seven *Citrus* species, including *C. australis* (an Oceanian species closely related to *C. australasica*) and *C. reticulata*. This anchoring was performed using the locOnRef tool from the Scaffhunter toolbox (Martin et al. 2016) allowing for the evaluation of synteny and collinearity between our assemblies and the reference map (Figure S1). The anchoring of each haplotype revealed a high degree of collinearity, with most chromosomes showing a clear one-to-one correspondence between physical and genetic coordinates. Minor discrepancies, visible as local discontinuities or small inversions in the Circos plots (Figure S1), likely reflect structural variants or orientation inconsistencies. These are expected when the available reference genomes differ from the actual parental genotypes used in the cross. Such local differences may also result from the trio-binning strategy, which preserves haplotypic divergence and can highlight micro-rearrangements between evolutionarily distant parental lineages. The absence of widespread inter-chromosomal misplacements and the consistent anchoring of large chromosome-scale scaffolds across linkage groups strongly support the structural accuracy of the haplotype-resolved assemblies. These results confirm that the assembly strategy, despite the unavailability of the actual parental genome sequences, produced high-quality haplotypes with conserved chromosome architecture and minimal technical artifacts.

The two haplotypes’ genomes are highly syntenic, although several structural variants can be detected as shown by a dot plot analyses with the presence of numerous structural variations, such as an inversion in the middle of the chromosome 1, a translocation on chromosome 3 and a large duplication on chromosome 5, for example (Figure 1B). Gene prediction *via BUSCO* analysis revealed high completeness for both haplotypes (98.4% for *C. reticulata* and 98.3% for *C. australasica*), with the majority of *BUSCO* present as unique or duplicated gene models (see Table 1).

**Table 1.** Summary statistics for the contigs of the hybrid diploid genome assembled with an haplotype resolution.

| Statistic | *C. reticulata* Haplotype | *C. australasica* Haplotype |
| --- | --- | --- |
| Genome size (Mb) | 349 | 359 |
| Number of contigs | 9 | 9 |
| N50 (Mb) | 38.0 | 39.9 |
| L50 | 4 | 4 |
| %GC | 34.13 | 34.10 |
| *BUSCO* completeness (%) | 98.4 | 98.3 |
| Unique genes (%) | 78.0 | 73.8 |
| Duplicated genes (%) | 19.7 | 24.5 |

### Deciphering the differential allelic expression in the inter-generic hybrid

Previous studies carried out on different combinations of hybrid plants have systematically revealed the presence of differential allelic expression to a greater or lesser extent within the hybrid compared with the parents, often for several hundred genes. To determine the situation within the hybrid between *C. australasica* and *C. reticulata*, we carried out a transcriptomic analysis in biological triplicate for the hybrid and each parent using leaf tissue. Within the hybrid, the transcripts were mapped onto the haplotype genomes in order to obtain an allelic resolution of the expression level of each of the genes.

To analyse differential gene expression between the two sub-genomes, we identified a common gene pool. Gene models were transferred from the reference *CITRE* (Droc et al. 2024) to each haplotype assembly using Liftoff (Shumate et Salzberg 2021). Liftoff is a stand-alone lift-over tool that takes as input the reference genome sequence (FASTA), the target genome sequence (FASTA), and the reference gene annotation (GFF/GTF) even if assembled genomes accumulate structural variation and divergence. It uses Minimap2 (H. Li 2018) to map each reference gene’s sequence onto the target assembly, and for each gene selects the set of exon alignments that maximizes sequence identity while preserving the exon–intron structure. Using this strategy, 95.0% of the (*CITRE*, ((Droc et al. 2024)) reference genes (33,055 out of 34,798) were successfully transferred to the maternal haplotype, and 89.0% (30,980 out of 34,798) were annotated on the paternal haplotype. Among these, 30,980 orthologous genes were found in both haplotypes and represent our pool of common genes.

We were thus able to determine which genes showed differential bias in transcript accumulation, and counted 1758 genes in which the contribution of the allele derived from *C. australasica* was at least 3 times greater, and 1809 genes in which the contribution of the allele derived from *C. reticulata* was 3 times greater within the hybrid (Figure 2A). Some transcript accumulation profiles were confirmed using RT-qPCR (Figure S2 and Table S3).

**Figure 2.**
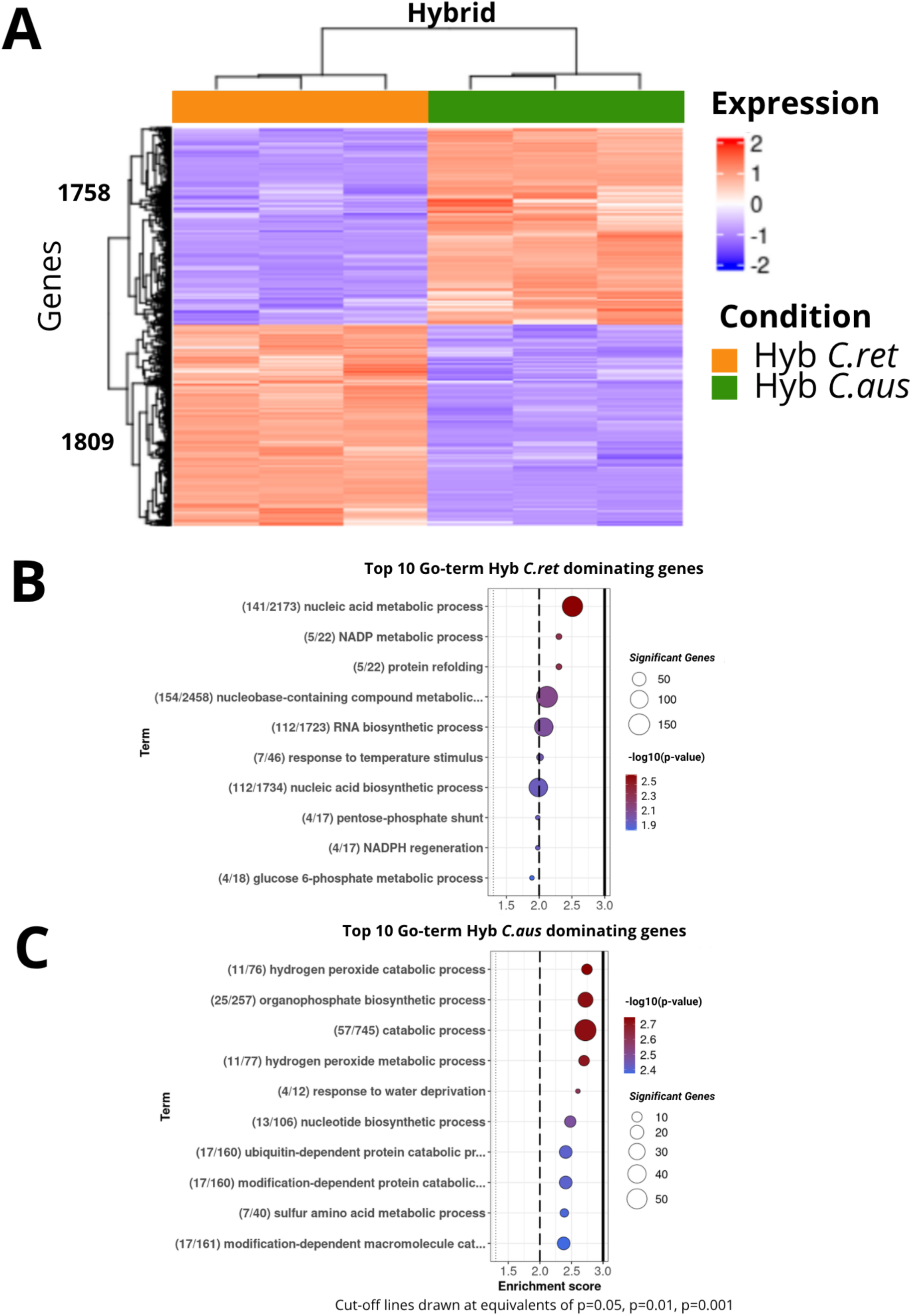
Allele-specific expression and functional enrichment in the hybrid. (**A**) Heatmap showing allele-specific expression between the two haplotypes in the hybrid. Expression values are variance-stabilized counts (VST) obtained from DESeq2 and centered gene-wise (z-score). Rows represent genes; columns represent biological replicates of the two hybrid haplotypes. The colour gradient reflects standardized expression levels: red indicates higher expression, blue indicates lower expression (unit: log₂-transformed normalized expression). (**B**) Bubble plot of enriched GO Biological Process terms among genes upregulated on the maternal haplotype. Bubble size represents gene count; colour reflects enrichment p-value. (**C**) Bubble plot of enriched GO terms among genes upregulated on the paternal haplotype, plotted as in panel (**B**).

We then determined whether certain classes of genes were particularly affected by this differential expression by determining changes in transcript accumulation according to gene ontology (GO term). Several groups of genes were found to be significantly differentially expressed within the hybrid. The *C. reticulata* haplotype is dominant for genes involved in metabolic processes for nucleic acid and NADP, while the *C. australasica* haplotype is dominant for genes implicated in catabolic processes particularly hydrogen peroxide, as well as in organophosphate biosynthetic processes (Figure 2B and 2C and Figure S3).

Such dominance at the GO term scale implies a biased implication of one of the parents in the hybrid in the regulation of certain metabolic and physiological pathways that could explain partially the phenotype observed in the hybrid.

Finally, we identified 289 and 268 genes in the hybrid that are exclusively expressed either by the *C. australasica* or the *C. reticulata* haplotypes respectively (Figure S4). Again, these numbers are very similar, demonstrating a somewhat overall balance between the two haplotypes. In both cases, the majority of these genes are only expressed in the corresponding parental lines (Figure S4).

### DNA methylation landscape in hybrid haplotypes reveals context- and parent-specific reprogramming

To determine whether some of this differential allelic expression may be the result of changes operating at the epigenetic level, we analysed cytosine methylation globally for the hybrid and each parent using leaf tissue. Whole-genome Bisulfite sequencing (WGBS) approaches were used and sequenced using 150bp paired-reads with depth of coverage equivalent to 20 times the average citrus genome size in biological triplicate.

Overall levels of DNA methylation revealed distinct patterns across the three cytosine contexts (Figure 3A). Methylated CG (mCG) were the most abundant, followed by methylated CHG and methylated CHH (mCHG and mCHH). Both hybrid haplotypes displayed a significant reduction in CG methylation relative to their respective parents, with Hyb *C. ret* showing a 5.2% decrease and Hyb *C. aus* a 5% decrease compared to *C. reticulata* and *C. australasica*, respectively. In contrast, mCHG levels were relatively stable across hybrids and parents. Notably, mCHH showed a haplotype-specific reprogramming: while Hyb *C. ret* maintained mCHH levels comparable to *C. reticulata*, Hyb *C. aus* exhibited a clear increase compared to *C. australasica*, suggesting targeted activation of RdDM in the paternal haplotype.

**Figure 3.**
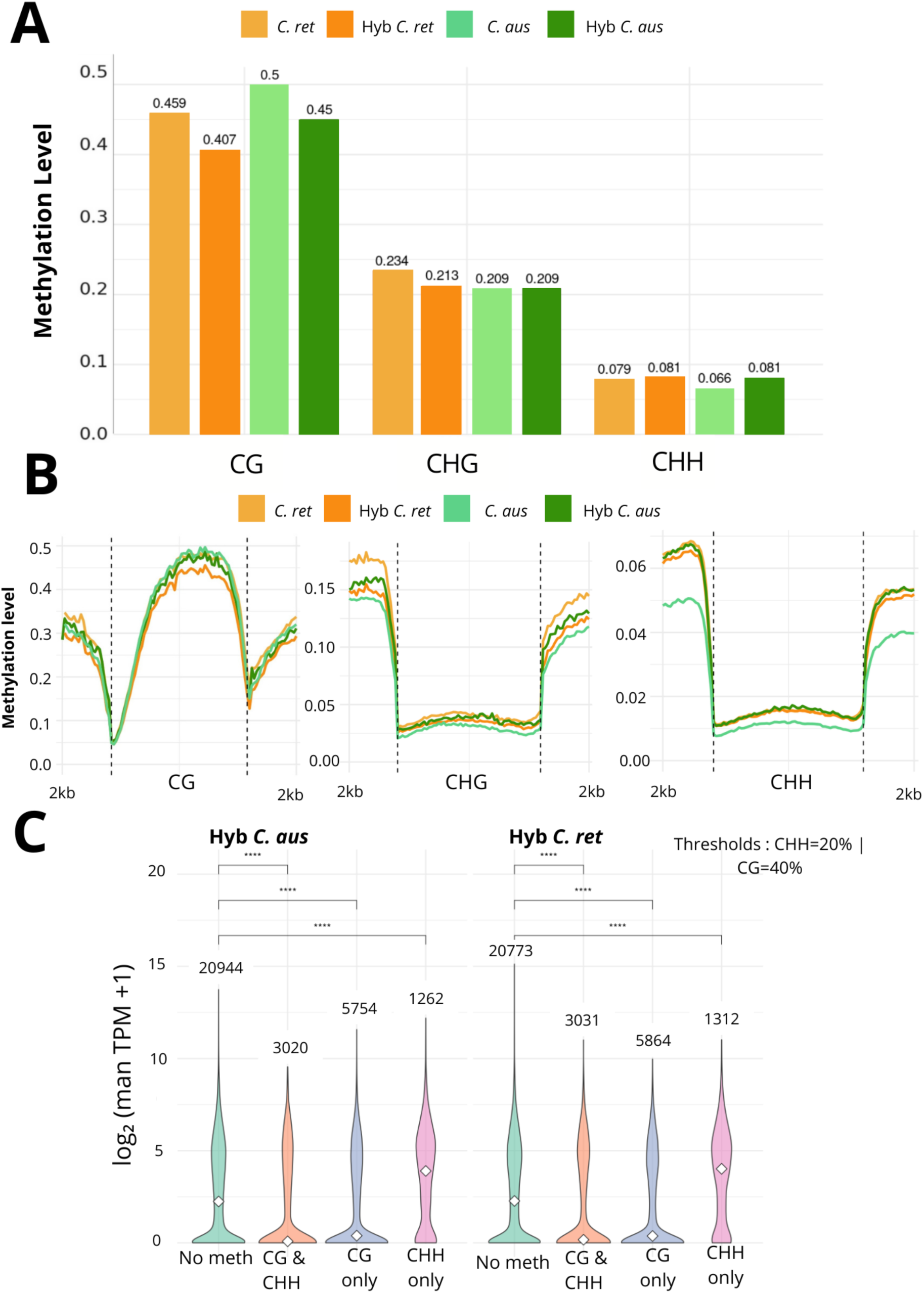
DNA methylation profiles and correlation with gene expression. **(A)** Global DNA methylation levels (in %) in CG, CHG, and CHH contexts across parental haplotypes (C. reticulata, C. australasica) and their corresponding haplotypes within the hybrid. Values indicate mean methylation levels across all annotated genes. **(B)** DNA methylation metaprofiles (in %) across gene bodies and ±2 kb flanking regions in each cytosine context. Curves represent average methylation profiles for each genotype and context. **(C)** Violin plots showing gene expression levels (log₂(mean TPM + 1)) according to promoter methylation status in CG and CHH contexts in the two haplotypes of the hybrid. Promoters are categorized into four classes: “No methylation”, “CG & CHH”, “CG only”, and “CHH only” based on thresholds (CHH ≥ 20%, CG ≥ 40%). Statistical differences in expression across categories were assessed using a Kruskal–Wallis test (p < 0.05), followed by pairwise Dunn’s post-hoc comparisons with Benjamini–Hochberg correction. The number of genes in each category is indicated above each violin. Asterisks denote significance levels from post-hoc comparisons: * = padj < 0.05, ** = padj < 0.01, *** = padj < 0.001, **** = padj < 1e-4.

At the chromosomal scale, genome-wide methylation distribution showed mCG and mCHG highest in pericentromeric regions, correlating with repetitive element abundance (Figure S5) (Giraud et al. 2025). Methylated CHH was generally lower but showed local peaks more pronounced in Hyb *C. aus* than Hyb *C. ret*, supporting reinforced RdDM activity targeting the *C. australasica* haplotype.

We then validated the methylation pattern observed using WGBS in the hybrid by analysing five different promoter regions that either accumulate or do not accumulate cytosine methylation using BS-PCR followed by Sanger sequencing (Figure S6 and table S3). Next, we analysed DNA methylation patterns across gene bodies and their flanking regions using metaplot analyses. In CHG and CHH contexts, methylation dropped sharply within gene bodies, with elevated levels in flanking regions (Figure 3B). mCHH around genes was markedly higher in *C. australasica* haplotype than in *C. reticulata* haplotype, reinforcing the idea of RdDM-driven reprogramming is more pronounced on the *C. australasica*-derived haplotype. Levels of mCG between the hybrid haplotypes were minor at the genic level, but a modest decrease in the *C. reticulata* haplotype was noted in promoter-proximal regions, whereas gene-body mCG remained relatively stable and high in both hybrids. These patterns reflect the complex locus- and context-specific DNA methylation reprogramming reported during cellular differentiation and hybridization ((Kakoulidou et Johannes 2024);(Lauss et al. 2018)) Direct comparisons between each hybrid haplotype and its respective parent confirmed an asymmetric pattern of methylation reprogramming (Figure 3B). Hyb *C. aus* showed increased mCHH compared to *C. australasica*, particularly in gene-flanking regions, while mCG was largely conserved. Hyb *C. ret* exhibited slightly elevated mCG relative to *C. reticulata*, with stable mCHH levels. This supports models where epigenetic remodelling in hybrids is haplotype-specific and influenced by parental origin, consistent with findings that DNA methylation remodelling in hybrids is often driven by epigenetic interactions and small RNA pathways (Kakoulidou et al. 2024).

We then looked at the impact of this methylation, whatever its context, on gene expression within the hybrid, with the idea of evaluating this type of epigenetic modification on the establishment of the differential allelic expression previously observed. For each haplotype, we analysed the relative expression of the genes when their promoter region (1 kb upstream of the transcription start site, TSS) was either unmethylated, or methylated at CG (at least 40%), or methylated at CHH (at least 20%), or both. As expected, genes whose promoters accumulate only mCG or both mCG and mCHH are strongly correlated with a low level of expression (Figure 3C), whatever the haplotype, compared to non-methylated promoters. In contrast, promoter regions only accumulating mCHH associate significantly with genes with a higher expression level than those without mCG only, both mCG and mCHH or in comparison with those that are unmethylated (Figure 3C). We also confirmed this result by analysing numerous genes where indeed high gene expression was not incompatible with the presence of small clusters and mCHH accumulation in their promoter (Figure S7). Taken together, our data reveal the presence of significant changes at the methylation levels, with contrast impact on gene expression depending on the cytosine context.

### Small RNA population and impact in the intergenic hybrid

In plants, mCHH accumulation in promoter regions is the result of the action of RdDM and RNA polymerases IV and V, which are specific to plants and enable the generation of small 24 nt RNA (hereafter named siRNA). Associated with an Argonaute protein (AGO4 in *A. thaliana*), these siRNA are then able to target sites for CHH methylation *via* the action of DRM2 (Matzke et al. 2015). Therefore, we expect that at least some of the promoters accumulating CHH methylation will also accumulate siRNA. To test this hypothesis, we performed small RNA-seq analyses for the hybrid and each parent using leaf tissue in triplicates.

Small RNA were grouped according to their size, showing the presence of two major types of small RNA: those of 21-nt, corresponding to microRNA, and those of 24-nt, which are more abundant and correspond to siRNA (Figure 4A) (Zhan et Meyers 2023). Overall, we can see that the two haplotypes participate equally in the production of these smallRNAs in quantitative terms (Figure 4B).

**Figure 4.**
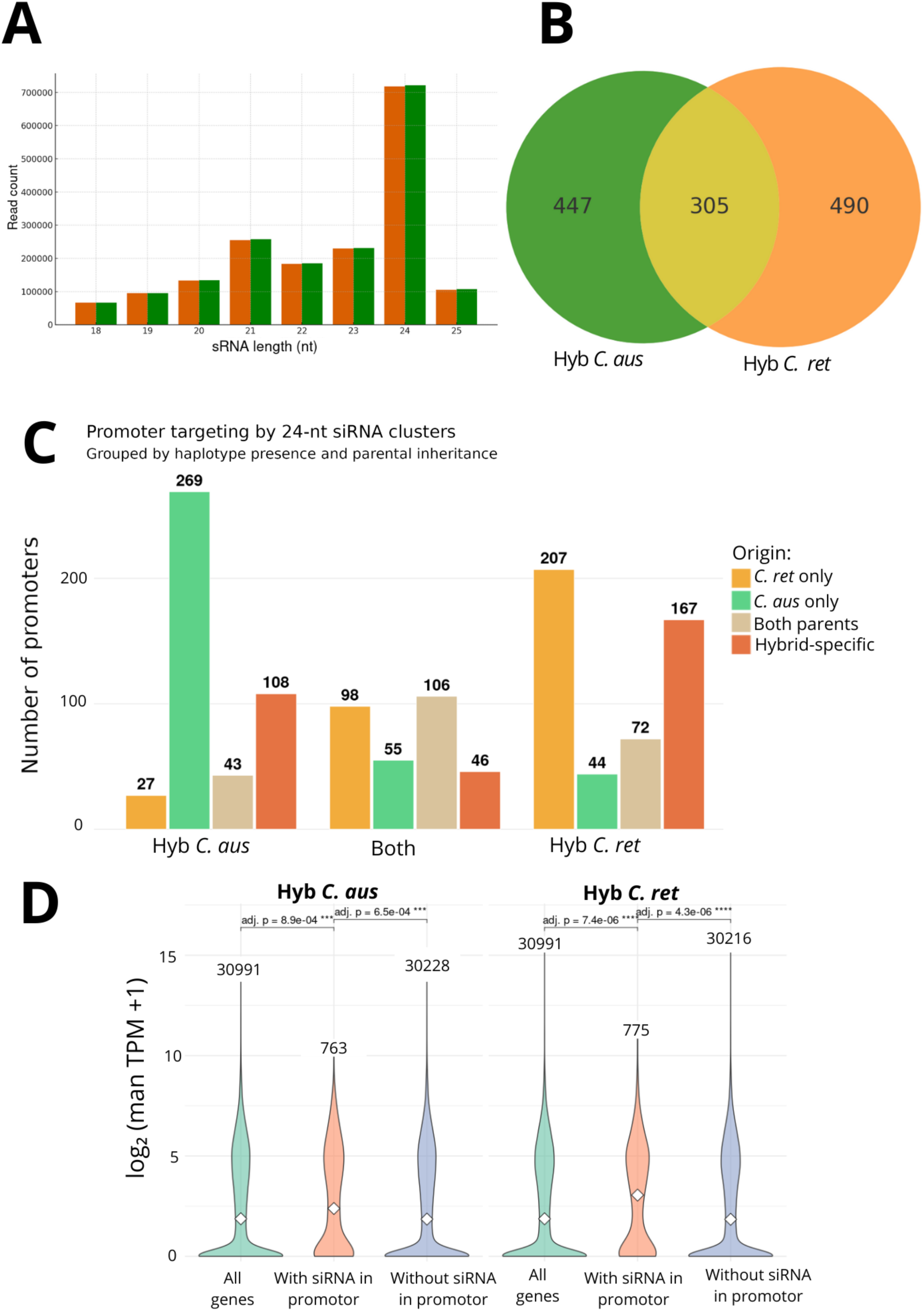
Haplotype-specific distribution and effects of 24-nt siRNA clusters. **(A)** Size distribution of mapped small RNAs in each hybrid haplotype**. (B)** Venn diagram showing the overlap of 24-nt siRNA clusters between the two haplotypes of the hybrid. The intersection represents clusters targeting the same promoters in both haplotypes. **(C)** Classification of gene promoters targeted by 24-nt siRNA clusters in the hybrid, based on parental origin. Promoters are grouped according to whether they are targeted by siRNA clusters inherited from C. reticulata only (orange), C. australasica only (green), both parents (beige), or are hybrid-specific (red). **(D)** Violin plots showing gene expression levels (log₂(TPM+1)) depending on the presence or absence of 24-nt siRNA clusters in promoters, for each hybrid haplotype. Genes with siRNA in their promoters tend to be more expressed than those without (Wilcoxon test, adjusted p-values indicated).

We then focused on the siRNA, responsible for targeting genomic regions in CHH methylation *via* the RdDM pathway. To analyse whether these siRNA target the same regions, we mapped them on the haplotype-resolved hybrid genomes. A third of the individual siRNA originates from both subgenomes, while the other two-thirds originate either from *C. reticulata* or *C. australasica*-derived haplotype (Figure S8). Because of the important sequence variation existing between the two haplotypes, we decided to question their functional impact once grouped into clusters when their coverage peaks were located within a 200-nucleotide window, as defined by ShortStack. Promoters targeted solely by CHH methylation are expected to be targeted by siRNA which then participate in the recruitment of DRM2 at promoter regions, then genes whose promoters are targeted by siRNA should also be more transcribed overall.

We therefore focused on the siRNA clusters targeting only the promoter regions (1 kb upstream of the TSS). Once again, the majority of siRNA groups were produced by only one of the two haplotypes within the hybrid. Only 25% were produced simultaneously by both haplotypes in the hybrid (Figure 4B). Among these haplotype-specific siRNA clusters, only half are also expressed in the parental lines (Figure 4C), highlighting a potential asymmetric inheritance. Finally, we analysed the global expression of the genes targeted or not by siRNA clusters in their promoter and demonstrated that those with targeted promoters accumulate more transcribed than those non-targeted in their promoter (Figure 4D). Altogether, these results demonstrate not only that the contribution of siRNA is essentially haplotype-dependent within the hybrid, but also that the targets of siRNA tend to be more transcribed, supporting the hypothesis of a potential involvement of RdDM in transcriptional activation within the hybrid.

### Impact of RdDM on differentially expressed genes (DEG) in the hybrid

To explore the impact of hybridization on the interrelation between gene expression, DNA methylation, and siRNA production, we integrated our epigenomic data at the haplotype level (Figure 5A–B). We then performed several statistical analyses of gene expression within the hybrid, quantified as TPM.

**Figure 5.**
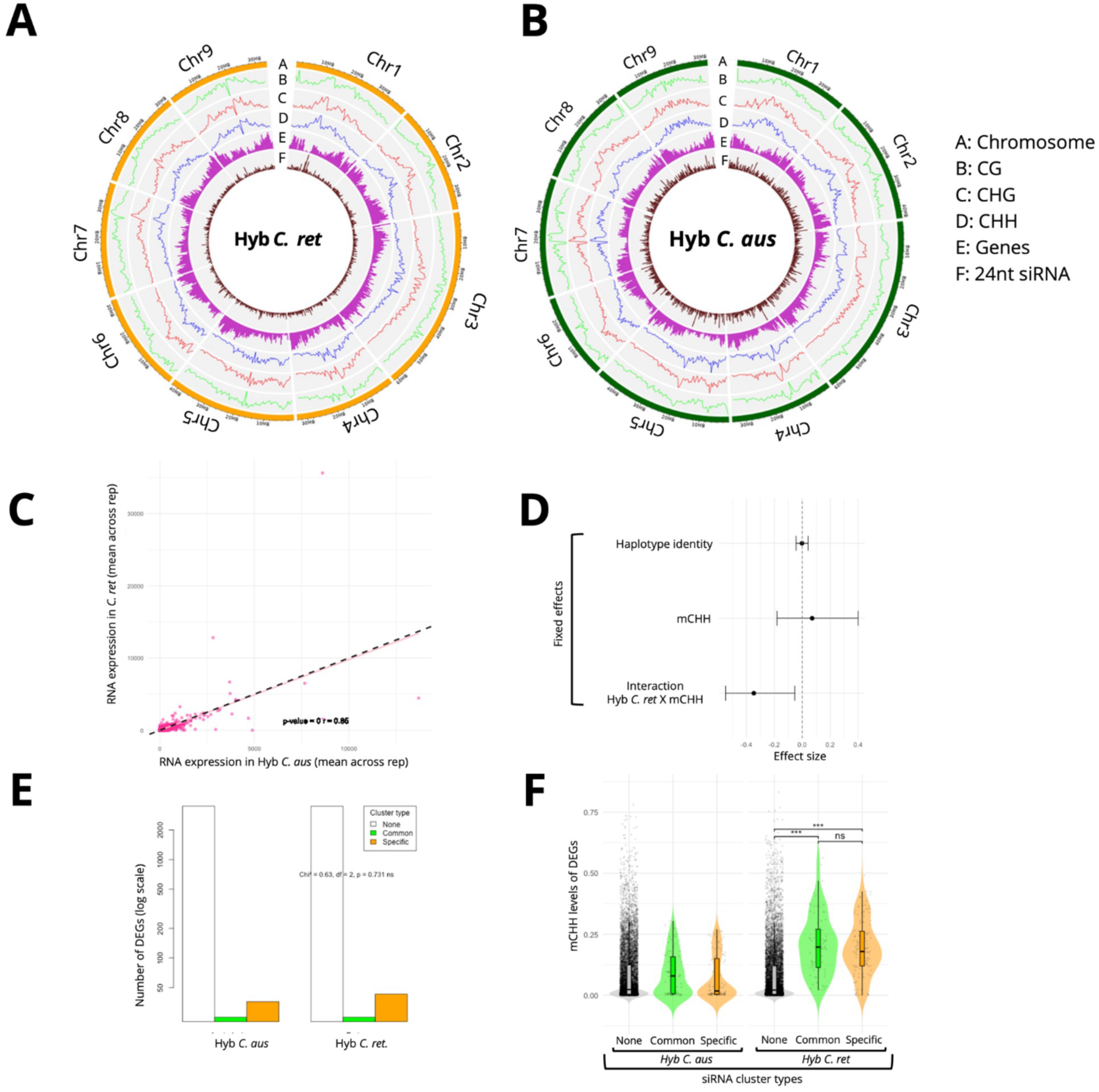
Genome-wide epigenomic patterns and overlap with expression. (**A–B**) Circos plots representing genome-wide profiles of DNA methylation, gene density, and 24-nt siRNA clusters across the nine chromosomes of the two haplotypes of the hybrid: (**A**) C. reticulata-derived haplotype (orange) (**B**) C. australasica-derived haplotype (green). Tracks from outer to inner rings: A : Chromosomes; B–D : DNA methylation levels in CG (red), CHG (blue), and CHH (purple) contexts; E : Gene density; F : Density of 24-nt siRNA clusters. (**C**) Spearman’s rank correlation in gene expression between haplotypes. (**D**) Effect sizes (±95% CI) from a linear mixed model evaluating the impact of haplotype, mCHH levels, and their interaction on DEG expression (TPM). Contrasts are treatment-coded with C. australasica haplotype as the reference. CIs overlapping zero indicate non-significance. (**E**) DEG distribution by siRNA cluster types and haplotypes (None = no cluster, Common = shared, Specific = haplotype-specific). (**F**) mCHH levels across small RNA cluster types by haplotype. Dunn’s test with Bonferroni correction: Significance: *** p ≤ 0.001, ** p ≤ 0.01, * p ≤ 0.05, ns = not significant.

First, we tested whether gene expression levels were correlated between haplotypes using Spearman’s rank correlation in R. We observed a strong and highly significant correlation (Spearman’s ρ = 0.85, *p* < 2.2 × 10⁻¹⁶), closely following a 1:1 relationship (Figure 5C), suggesting that most genes exhibit similar expression levels across the two haplotypes. We then tested whether the 3,567 DEG within the hybrid were influenced by haplotype identity, mCHH levels, or the interaction between the two. We used a linear mixed-effects model with haplotype identity, mCHH and the interaction included as fixed effects, while the first principal component (PC1) from a PCA of gene expression profiles was included as a random effect. This approach captures underlying clustering or co-expression patterns between genes that could confound the interpretation of haplotype and methylation effects. The model revealed no overall differences in gene expression of DEG between haplotypes, confirming that neither haplotype systematically dominates the other in terms of expression of DEG (Top; Figure 5D). It also showed that mCHH alone had no consistent effect on gene expression on DEG (Middle; Figure 5D). However, a significant interaction between haplotype identity and mCHH levels was detected (Bottom; Figure 5D): a marked reduction in gene expression was specifically observed when mCHH occurred on the *C. reticulata* haplotype compared to *C. australasica* haplotype. This suggests that the effect of mCHH on gene expression of DEG is haplotype-dependent.

We then investigated the impact of hybridization on siRNA clusters on promoters of DEG. We categorised clusters in three classes: “Specific” (detected in only one haplotype), “Common” (shared between both), and “None” (genes with no detected cluster in their promoter). We counted the number of DEG per category and haplotype, and used a Pearson’s Chi-squared test of independence to assess differences in distribution. The distribution of siRNA cluster types did not significantly differ between haplotypes (Chi² = 0.63, df = 2, *p* = 0.73). Most DEG lacked a detectable cluster in both haplotypes, and the proportions of “Common” and “Specific*”* clusters were similar across haplotypes (Figure 5E). These results suggest that the presence or specificity of siRNA clusters among differentially expressed genes is similar across haplotypes.

Finally, to test whether mCHH levels differed on promoter regions targeted by siRNA clusters according to their origin, we performed another test, still focusing on DEG. Since methylation data did not follow a normal distribution, we used non-parametric Kruskal-Wallis rank sum tests to compare methylation levels across cluster types (“None”, “Specific”, “Common”) separately for each haplotype. In the *C. australasica* haplotype, mCHH levels did not differ significantly between cluster types (*χ²*(2) = 2.35, *p* = 0.31). No pairwise comparisons were significant after correction. In contrast, *C. reticulata* haplotype showed a significant effect of siRNA cluster type on mCHH (*χ²*(2) = 232.25, *p* < 2.2×10⁻¹⁶). Post hoc tests revealed that DEG with no associated siRNA cluster had significantly lower methylation than those with either “common” or “specific” clusters (*p* < 0.001 for both comparisons). Moreover, there was no significant difference between “common” and “specific” clusters (*p* = 0.67 after Bonferroni correction). These results suggest that in *C. reticulata* haplotype, the presence of a siRNA cluster regardless of whether it is shared or specific is associated with an increase in mCHH, whereas in *C. australasica* haplotype, mCHH is not significantly affected by cluster occurrence (Figure 5F).

### Association of RdDM with expressed genes is not linked to hybridization

Our strategy revealed that while for the majority of genes, the parental contribution to transcript accumulation within the hybrid was balanced, around 10% of common genes were dominated by the contribution of just one of the two parents. This dominance is even more marked for siRNA, where only a third are expressed by both haplotypes.

Genome-wide analyses showed that cytosine methylation, across CG, CHG, and CHH contexts, accumulates in gene-poor regions near the chromosomal centres, likely corresponding to centromeric or pericentromeric regions, despite centromeres not being clearly annotated in Citrus (Xia et al. 2020). Both haplotypes display consistent patterns of mCHH and siRNA accumulation in these regions, indicative of canonical RdDM activity, potentially targeting transposable elements. The strong co-localization of mCHH and siRNA in pericentromeric regions further supports their coordinated role in heterochromatin maintenance, although siRNA-independent mCHH accumulation may also be involved ((Stroud et al. 2014) ; (He et al. 2021)). Despite the interspecies nature of the hybrid (*C. reticulata* × *C. australasica*), the epigenomic landscapes of both haplotypes show remarkable global symmetry. This conserved chromosomal organization suggests that genome compartmentalization and RdDM activity are stable features across divergent Citrus lineages and may contribute to the overall transcriptional stability observed in the hybrid (Figure 5A–B), although a potential, we identified a haplotype-bias impact in the hybrid for DEG (Figure 5F).

To assess whether RdDM is linked to transcriptional activity, we analysed gene expression levels in relation to both mCHH accumulation and siRNA presence. As previously shown, analysis of expression data with either cytosine methylation or the presence of siRNA revealed that genes were more highly expressed when they either accumulated mCHH (Figure 3) or were targeted by siRNA (Figure 4). To see if these two aspects are linked, we analysed all the genes targeted by siRNA and that also accumulate at least 20% mCHH in their promoter region, separating the expressed genes (Figure 6A) from the genes that are little or no expressed (Figure 6B). Remarkably, genes whose promoters accumulate these two characteristics are 10 times more likely to be associated with expressed than unexpressed genes. However, these targeted genes are only targeted within the hybrid in at least half the cases: in 49% of cases (93 promoters out of 191) when the genes are associated with expression (Figure 6A) and in 77% of cases (24 promoters out of 31) when the genes are not or only slightly expressed (Figure 6B).

**Figure 6.**
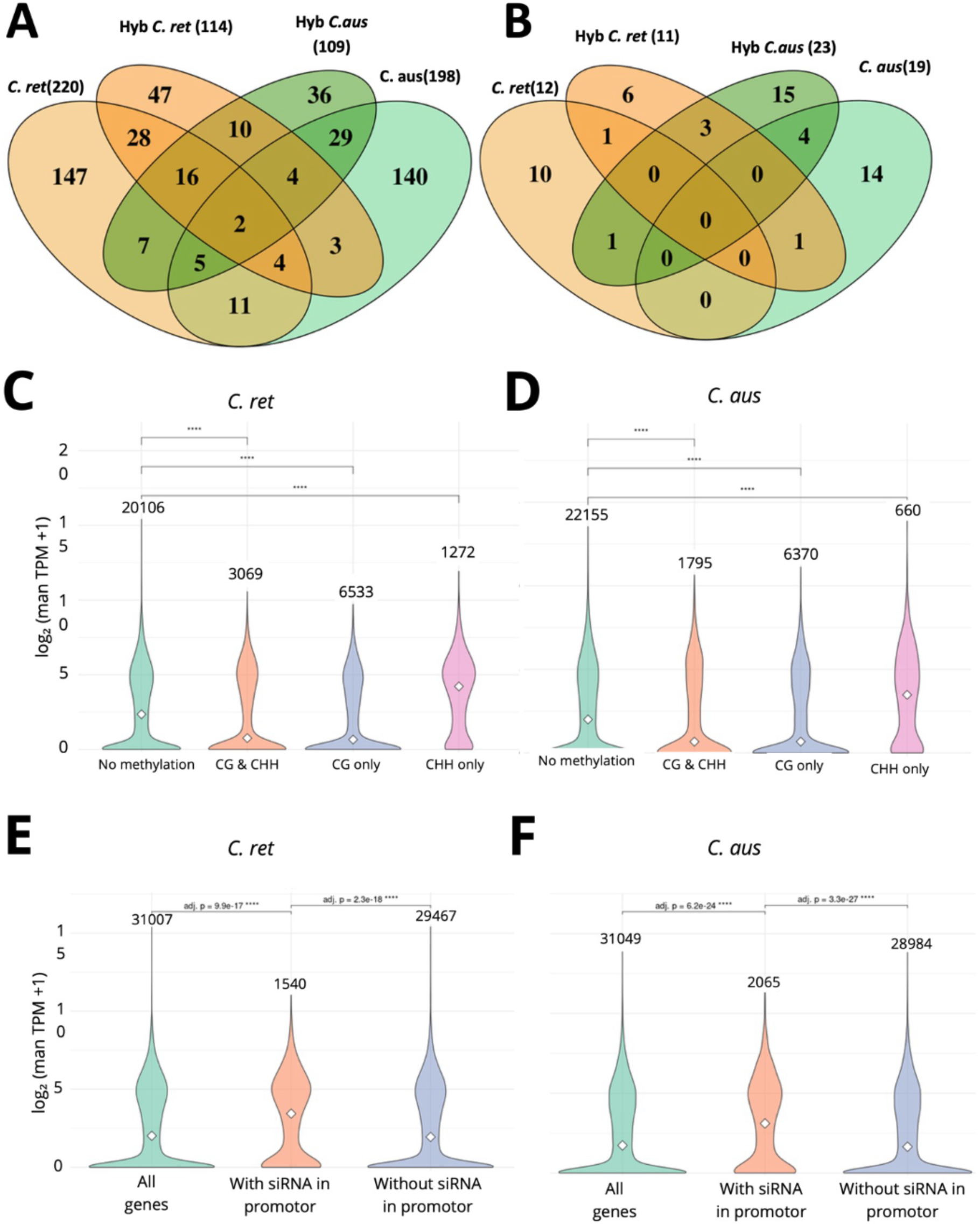
Epigenetic regulation of gene expression in the parental genomes. (**A–B**) Venn diagrams showing the overlap of genes whose promoters are targeted by 24-nt siRNA clusters across parental and hybrid haplotypes. (**A**) Expressed genes (TPM > 1) with promoter siRNA clusters. (**B**) Non-expressed genes (TPM ≤ 1) with promoter siRNA clusters. Each subset indicates the number of genes whose promoters are targeted by siRNA in one or more haplotypes**. (C–D)** Violin plots showing gene expression levels (log₂(mean TPM + 1)) in parental genomes: **(C)** C. reticulata, **(D)** C. australasica. Genes are grouped according to promoter methylation status: No methylation, CG only, CHH only, and CG & CHH, using thresholds of 40 % CG and 20 % CHH methylation. Numbers above each plot indicate the number of genes in each category. Asterisks indicate significant differences (Wilcoxon test, **** = p < 1e-4). **(E–F)** Gene expression comparison based on the presence or absence of 24 nt siRNA clusters in promoters for: **(E)** C. reticulata, **(F)** C. australasica. Wilcoxon rank-sum tests were used to assess statistical differences; adjusted p-values are indicated (Benjamini–Hochberg correction).

To test whether RdDM targets are also associating with higher expression in the parental lines, we analysed the transcriptomic, DNA methylation and small RNA data in the parents, using the haplotype resolved reference genome of the hybrid (Figure S9). Interestingly, mCHH is also linked to high gene expression in the parental lines, which goes to show that this atypical correlation is not linked to hybridization *per se*, but would rather be a more general feature in *Citrus* genus. A closer examination of siRNA–CHH methylation modules shows that both CHH methylation and siRNA accumulate specifically within the 1 kb region upstream of the transcription start site (TSS), mostly at expressed genes (Figure S10).

To confirm this hypothesis, we perform the same analyses made on the hybrid haplotype individually on methylation and siRNA accumulation in the parental lines, and indeed confirm a similar association of expressed genes with siRNA cluster or with only mCHH both in *C. reticulata* (Figure 6C and E) and in *C. australasica* (Figure 6D and F). *In fine*, in both *C. reticulata* and *C. australasica*, genes with promoter accumulating mCHH and/or siRNA clusters were more likely to be expressed than genes without these marks, reinforcing the conclusion that this epigenetic pattern is conserved across species. Venn diagram analysis further showed partial overlap between mCHH and siRNA-targeted genes in each parent (Figure S11), suggesting that these two layers of RdDM regulation may act independently or additively.

To assess whether this profile is shared across citrus species, we took advantage of the published RNA-seq and WGBS-seq data in citrus lemon (*Yu H. et al., 2024).* Again, a significant higher expression is also observed for the 783 genes that accumulate CHH methylation in their promoter (Figure 7, left panel). Moreover, as only leaf tissue has been analysed so far, we also analysed the correlation between gene expression and accumulation of CHH methylation only in the promoter for other citrus and related species, this time using root tissues (Figure 7). WGBS and RNA-seq analyses, from *C. reshni* (Cleopatra mandarin, a member of *C. reticulata*) and *Poncirus trifoliata,* were performed for this study. In addition, we also re-analysed advantage of the methylation and expression data previously published for *C. sinensis* (Huang et al., 2021). In these three species, we were able to confirm a significant increase in gene expression when CHH-only methylation accumulates in gene promoters (1 kb upstream of the TSS) (Figure 7).

**Figure 7:**
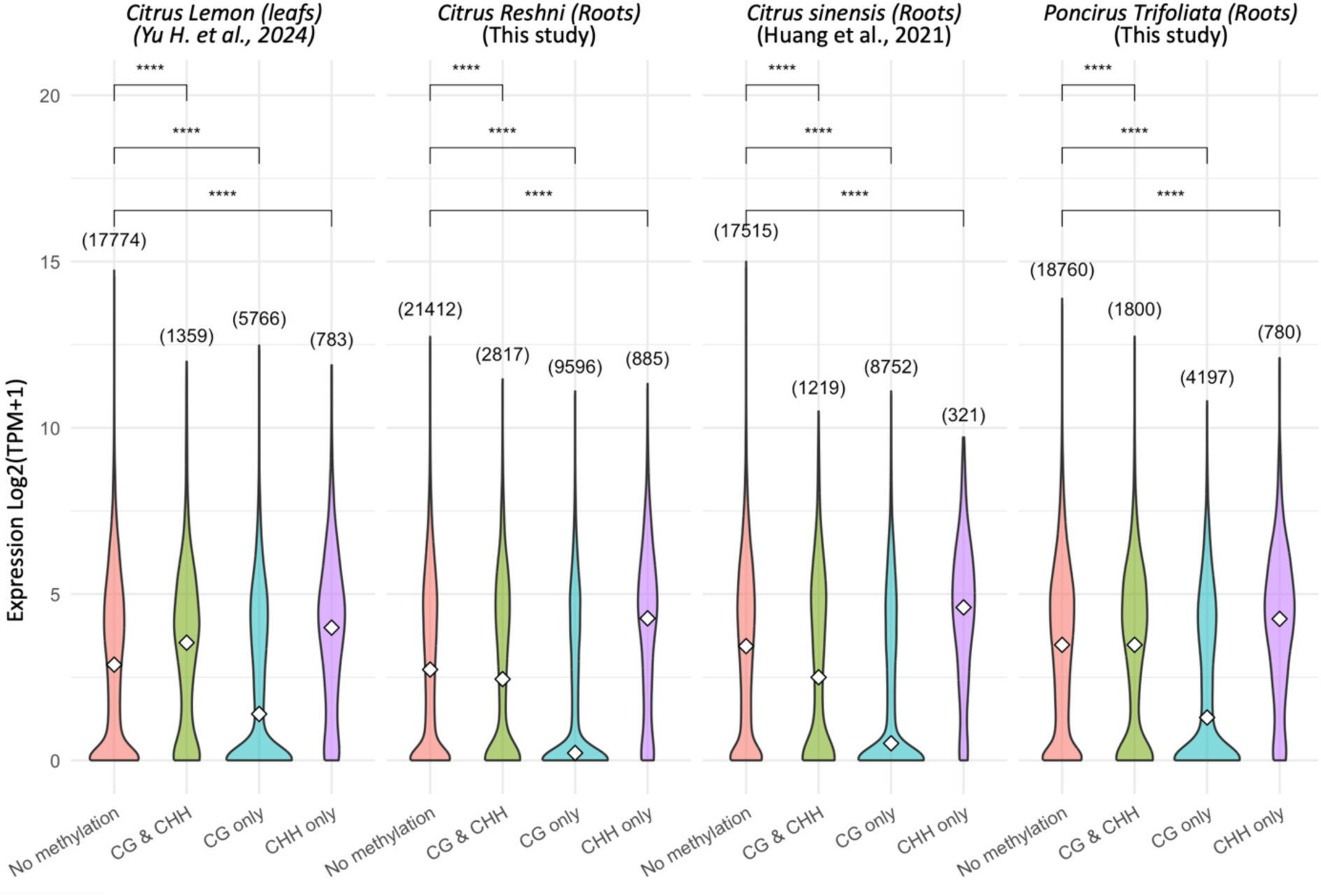
Correlation between DNA methylation and expression profiles in root and shoot tissues for Citrus and related species. Violin plots showing gene expression levels (log₂(mean TPM + 1)) according to promoter methylation status in CG and CHH contexts in Citrus lemon, Citrus Reshni, Citrus sinensis and Poncirus trifoliata. Promoters are categorized into four classes: “No methylation”, “CG & CHH”, “CG only”, and “CHH only” based on thresholds (CHH ≥ 20%, CG ≥ 40%). Statistical differences in expression across categories were assessed using a Kruskal–Wallis test (p < 0.05), followed by pairwise Dunn’s post-hoc comparisons with Benjamini–Hochberg correction. The number of genes in each category is indicated above each violin. Asterisks denote significance levels from post-hoc comparisons: **** = padj < 1e-4.

Altogether, these results support a model in which RdDM-associated targeted genes are positively associated with gene expression in *Citrus*, and this association is conserved across divergent species. If the results suggest stable associations across datasets, they do not strictly exclude the influence of stage- or environment-specific factors.

## DISCUSSION

In this study, we investigated the transcriptional and epigenetic consequences of an interspecific hybridization event between two phylogenetically distant *Citrus* species separated by over 4 million years of divergence: *C. reticulata* and *C. australasica*. By integrating high-quality haplotype-resolved genome assemblies with transcriptome, DNA methylation, and small RNA datasets, we provide new insights into the regulatory dynamics occurring in such distant hybrid. To analyse the citrus hybrid and its parent, we developed a pipeline adapted to the integration of genomic and epigenomic data with a haplotype resolution using the strategy previously published in (Koren et al. 2018) and that resolves allelic variation before assembly. This trio-binning approach enabled the separation of two highly divergent haplotypes and produced chromosome-scale assemblies with high contiguity and collinearity (Figure1 and S1). The observed structural variants, including small inversions and local discontinuities, may reflect true polymorphisms between the actual parental genomes or arise from hybrid-specific chromosomal rearrangements. While the assemblies already provide a robust foundation for genomic comparisons, future polishing steps, for example through the integration of Hi-C data, could further improve scaffold orientation and help distinguish between biological rearrangements and minor assembly inconsistencies. More importantly, the haplotype-resolved assembly provided the basis for identifying differentially expressed genes with allelic resolution in the hybrid. Because all analyses were conducted under non-stressed conditions, in the absence of both biotic and abiotic challenges, the regulatory patterns described here should be interpreted as a basal, haplotype-resolved framework rather than as direct evidence of stress- or disease-resistance mechanisms.

Hundreds of expressed genes accumulate transcripts mainly from only one of the parental-derived genomes, which implicate a certain dominance at the gene scale (Figure 2, S3 and S4). Although some metabolic or catabolic pathways are enriched among the dominant genes, we do not know at this stage the impact of this imbalanced expression. However, we can note that this imbalance seems to be somehow equilibrated between the two parent-derived genomes since a similar number of DEG are found in the haplotype-resolved hybrid transcriptome.

At the methylation level, the global epigenomic landscapes were remarkably conserved between haplotypes, closer inspection of Figure 5A–B reveals subtle but potentially meaningful differences in local siRNA accumulation. Discrete siRNA peaks specific to one haplotype, particularly on chromosomes 6 and 9, may reflect species-specific bursts of transposable element (TE) activity or structural rearrangements. These features could mark regions of differential genome organization between *C. reticulata* and *C. australasica,* but definitive interpretation will require high-quality TE annotation of each phased haplotype.

Quantitatively, if a global decrease in mCG and mCHG is observed in the hybrid compared to the parent, mCHH is slightly higher, suggesting a particular role or deregulation of the RdDM pathway. Interestingly, mCHH only significantly affects DEG derived from the *C. reticulata* haplotype (Figure 5D and S12). This haplotype-specific mCHH methylation is accompanied by a specific accumulation of siRNA in the *C. reticulata* haplotype promoter of DEG (Figure 5F).

The reason why only one haplotype seems to be sensitive to RdDM is not clear. mCHH accumulation in the hybrid seems to be additive, reaching the highest level of the parent as shown in (Figure 3B). In consequence, we detected an increase in the level of mCHH of around 50% throughout the genome of the *C. australasica* haplotype compared to the parental line. This increase, in one generation, is considerable and could probably explain at least partially why mCHH do not have as much impact on gene transcriptional regulation in the *C. australasica* haplotype than in the *C. reticulata* haplotype, within the hybrid. One explanation could be that because of the sequence variation between the two haplotypes, mCHH readers may act preferentially on their own haplotype genome and a 50% increase is probably due to deregulation and/or an imbalance that cannot be corrected in this generation, saturating mCHH reader belonging to the *C. reticulata* haplotype. In addition, we cannot entirely rule out the potential impact of stage- or environment-specific factors. Previous studies in *A. thaliana* hybrids demonstrated that altered DNA methylation and siRNA patterns exist in the hybrid, potentially contributing to the hybrid vigor phenotype (Greaves et al. 2015). However, in that case, an increased mCG and a decreased mCHH has been observed relative to the parents (Greaves et al. 2012), which is the reverse of our observation here. Further transcriptional and epigenetic analyses on future generations will show whether this imbalance persists.

We suggest that the RdDM pathway is much more implicated in transcriptional activation, than repression in *Citrus*. One hypothesis could be that the RdDM pathway is affected in citrus, with transcriptional regulation affecting DNA methylation deposition or maintenance. However, the expression profile of the major members of the pathway appears stable in the hybrid and parents (Figure S13). As a fact, we identified hundreds of highly expressed genes in the hybrid, but also in the parental lines that accumulate both mCHH to a level higher than 20%, and that are also targeted by siRNA at their promoter. This number of targets is probably higher since it cannot be ruled out that the lack of detection of siRNA clusters is not the result of transcriptional variation due to the transient nature of their production. These characteristics are typical of RdDM targets. In contrast, only a few low-expressed genes have at least one of these characteristics in the three different genomic citrus contexts. Generally, RdDM is strongly associated with gene transcriptional repression in *A. thaliana*, maize, tomato and other plant species ((Rymen et al. 2020);(Law et Jacobsen 2010)). However, such a paradigm has been recently challenged by recent studies. In rice, the presence of mCHH on transgenes and endogenous sequences is not always linked to gene repression (Okano et al. 2008). In *Poncirus trifoliata*, a close relative of the citrus genus usually used as rootstock, highly expressed genes were more likely to accumulate mCHH sequence contexts in their promoter regions in this species (Wang et al. 2022). The molecular mechanisms that explain the potential involvement of RdDM either in transcriptional repression or activation have been revealed in *A. thaliana*, *via* the identification of different readers of methylated CHH that belong to SUVH histone methyltransferases (Pontvianne et al. 2010). When mCHH is associated with repression, SUVH4, 5 and 6 are recruited. But when mCHH is associated with high expression, SUVH1 and SUVH3 are implicated instead (Harris et al. 2018). These homologs also exist in citrus and are expressed in the parental and the hybrid plants. Creating SUVH1 and SUVH3 methylation reader knock-out using CRISPR-cas9 Should reveal, in the future, if the association is mediated by these specific histone methyltransferases. Furthermore, we could investigate the impact of CHH methylation on highly expressed genes by generating knockouts of essential RdDM core components, such as the *de novo* methyltransferase DRM2 or the catalytic subunit of RNA polymerase V. One additional possibility is that open chromatin states and high transcription may favor de novo CHH methylation, as previously observed in chromatin mutants in *Arabidopsis thaliana* (Earley et al. 2010). This hypothesis could be tested by changing the chromatin status of different identified promoter using methylase or demethylase and see how it affects siRNA accumulation and gene expression (Gallego-Bartholomé J. 2020). However, while CRISPR-Cas9 technology is feasible and established protocols exist in citrus (Wang et al. 2023), it remains technically challenging and time-intensive. As an alternative approach, we propose to investigate the relationship between CHH methylation patterns and gene expression in citrus under stress conditions or across different developmental stages. This will allow us to assess how these genes respond when their methylation or expression profiles are experimentally altered.

## CONCLUSIONS

Our study reveals the impact of DNA methylation and siRNA on the establishment of DEG in a complex inter-specific citrus hybrid. However, additional mechanisms can explain the presence of so many DEG in the hybrid. For example, structural variation including duplication, translocation and inversion contribute to the generated genomic diversity and gene copy number changes that can lead to phenotypic plasticity, including in crops (Picart-Picolo et al. 2020). However, their accurate detection requires a near perfect assembly, usually assisted with chromatin-chromatin interaction maps (Hi-C) approaches which is not our case here.

Mobile genetic elements such as transposable elements (TE) can also affect gene expression in two ways: either *via* creating structural variation *via* their neo-insertion, or by modulating gene expression in their vicinity ((Emmerson et Catoni 2025),(Zhang et al. 2023), (Thieme et al. 2022) (Baduel et al. 2019)). Their presence in promoter regions can therefore impact siRNA production and inheritance in hybrid context (Y. Li et al. 2012) . However, in addition to a near perfect assembly, a fine-tuned TE annotation is required in that case and could be the subject of a future study on this hybrid. Finally, mono-allelic expression of certain genes in this complex hybrid may be the consequences of the evolutionary process and the accumulation of genetic and epigenetic modifications in the promoter regions, affecting transcription factors association. Indeed, distant by 4 million years of evolution, the co-evolutionary process between the transcription factors and their targeted regulatory elements in gene promoters may not be fully conserved between the two species.

In any case, this study proposes an innovative and effective analysis strategy for analysing the molecular consequences of hybridization between distantly related species within the citrus genus. We hope that this strategy will be used in the near future to demonstrate (*i*) the reproducibility of DEG profiles and the impact of DNA methylation and siRNA production in several independent F1 hybrids between these two distantly related species, *C. reticulata* and *C. australasica*, and (*ii*) the ability to combine genomic and epigenomic approaches on young citrus hybrids to anticipate their phenotype during their reproductive stage in terms of their ability to resist pathogens such as Huanglongbing (Graham et al. 2024), but also to produce fruit that meets the standards for marketing.

## MATERIALS AND METHODS

### Plant material

Hybrid F1 result from a cross between *Citrus reticulata* Blanco cv Fortune mandarin (female parent) and *Citrus australasica* cv Caviar Bachès (Male parent), located at the INRAE CIRAD station in San Giuliano, France. The tissue analysed for *P. trifoliata* and *C. reshni* has been previously described in (Bonnin et al. 2024). The plant material used to analyse *Citrus lemon* and *Citrus sinensis* has been previously described in (Yu et al, 2024) and (Huang et al, 2021) respectively.

### Long-Read Sequencing

Genomic DNA from the hybrid was sequenced using the Oxford Nanopore Technologies (ONT) PromethION platform at the LGDP sequencing facility in Perpignan, generating a total of 63.88 Gb of long-read data. This corresponds to approximately 182X coverage of the ∼350 Mb genome.

### Genome Assembly and Annotation

Nanopore reads from the hybrid were assigned to parental haplotypes using a trio-binning approach. K-mers specific to each parent were extracted from Tell-Seq data using Meryl. Reads were assembled separately for each haplotype using Canu v2.2 (Koren et al. 2017) with options *genomeSize=430m*. Initial polishing of the assemblies was performed using Medaka (Wick 2016), followed by a second round with Pilon (Walker 2014) using the mapped Tell-Seq reads and the options *--fix snps,indels --minmq 3*.

### Gene annotation transfer

Gene models were transferred from the reference genomes using Liftoff (Shumate et Salzberg 2021) with the parameters *--copies --polish --exclude_partial,* ensuring accurate mapping of complete annotations.

Additionnal supporting information can be found in the supplemental material and methods document.

## ACKNOWLEDGMENTS

The authors would like to thank Raphaël Morillon, Patrick Ollitrault, François Luro, Anne Roulin, Flavia Thiebaut, Marie-Christine Carpentier, Moaine Elbaidouri, Gaëtan Droc, and Gauthier Sarah for fruitful discussions. This work received support from the sequencing and Bioinformatic platforms “Bio-environnement” at the University of Perpignan, the MESO@LR-Platform at the University of Montpellier and technical support from the bioinformatics group of the UMR AGAP Institute, member of the South Green Bioinformatics Platform. We would like to thank the “Occitanie Region” for funding KD and the “Collectivité de Corse” for funding MB. The authors are also grateful to Michel and Benedicte Bachès for sharing leaves to analyse the parental line *C. australasica*.

## CONTRIBUTIONS

A. G. performed the statistical modelling presented in FIG 5C-F. Y. F. created the citrus hybrid. K. D. extracted the RNA and DNA from the parent and the hybrid with the help of N. P. and F.P. M.B. realised RNA-seq and WGBS analyses for *C. reshni* and *P. trifoliata*. C.L. performed nanopore sequencing. F.P., N.P. and B. H. conceived and designed all experiments. K. D. performed all bioinformatic analyses with some help of M-C.C and N.P. and B.H. F.P. wrote the article with substantial contributions of K.D., B.H and A. G.. N.P. edited the article. N.P. and F.P. acquired main funding. F.P., N.P. and B.H. coordinated the research. All authors approved submission. All authors have read and agreed to the published version of the manuscript.

## DATA AVAILABILITY STATEMENT

Nucleotide sequencing data generated in this study have been deposited in European Nucleotide Archive (ENA) under the accession number PRJEB94133 (https://www.ebi.ac.uk/ena/browser/search/PRJEB94133).

## Supplemental Figures and Tables

**Figure S1.**
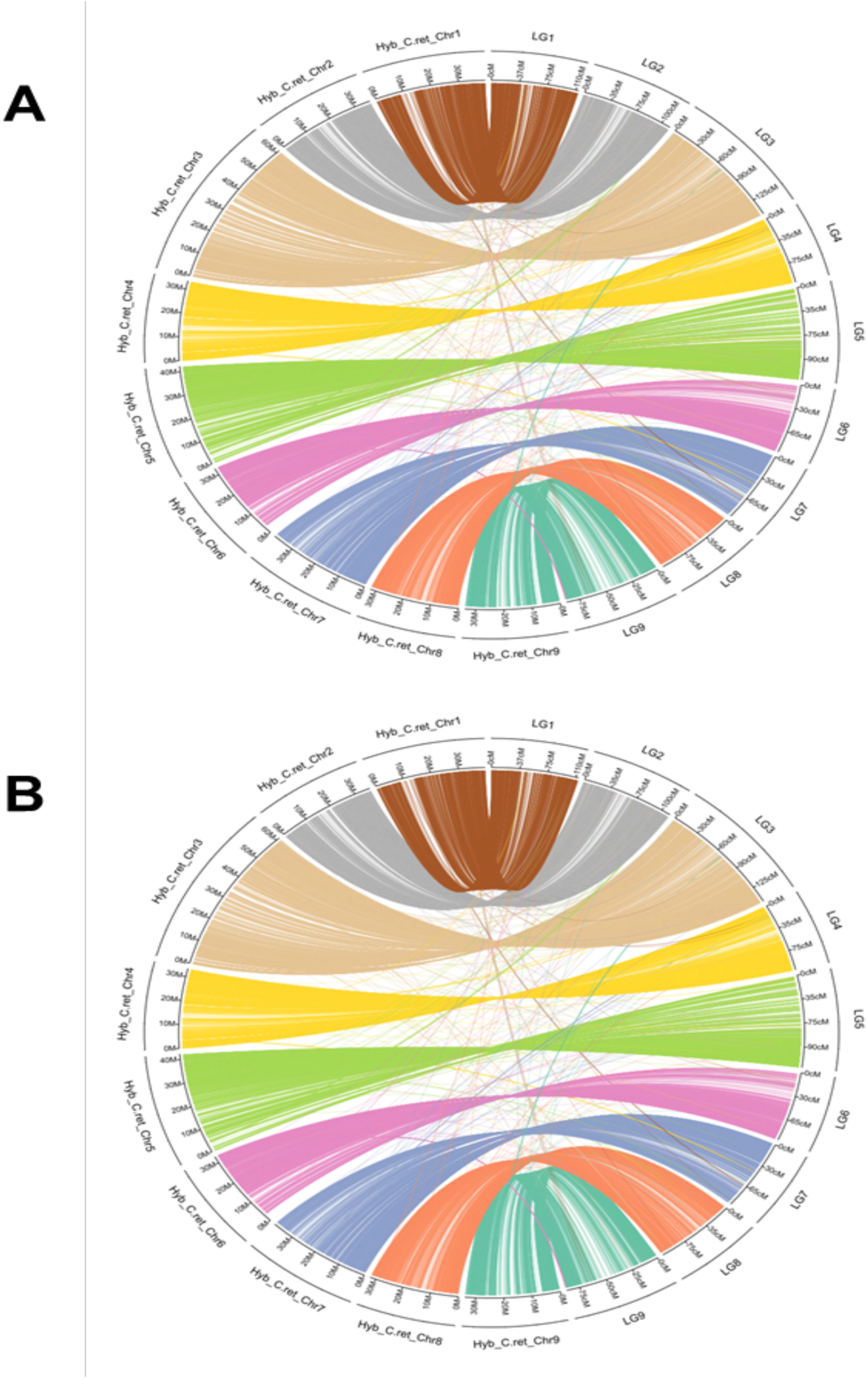
Anchoring of the haplotype assemblies to the consensus Citrus genetic map. Each ribbon connects a genomic scaffold from the haplotype assembly (left) to its corresponding position in one of the nine linkage groups (right). (**A**) Haplotype derived from C. reticulata. (**B**) Haplotype derived from C. australasica. The strong one-to-one correspondence and limited cross-linking support the structural accuracy of the assemblies.

**Figure S2.**
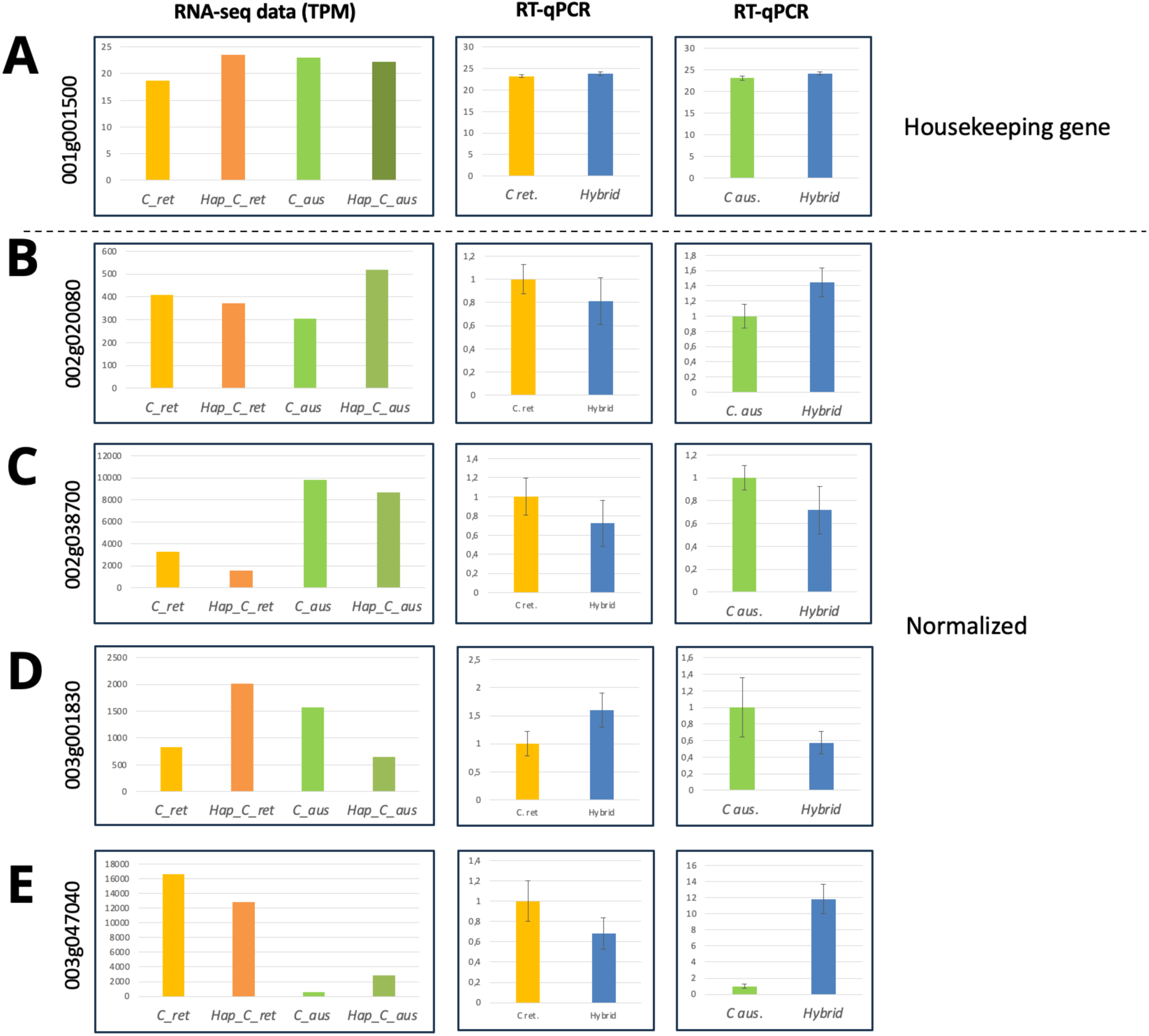
Validation of RNA-seq analyses by RT-qPCR. Analyses of transcripts accumulation of five different expressed genes in the parental and the hybrid lines according to the RNA-seq analyses (left graphic) or by RT-qPCR using C. reticulata-specific primers (middle praphic) or C. australasica-specific primers (right graphics). RT-qPCR of genes presented in the panel **B-D** were normalized according the housekeeping gene encoding UBIQUITIN (panel **A**; 001g001500).

**Figure S3.**
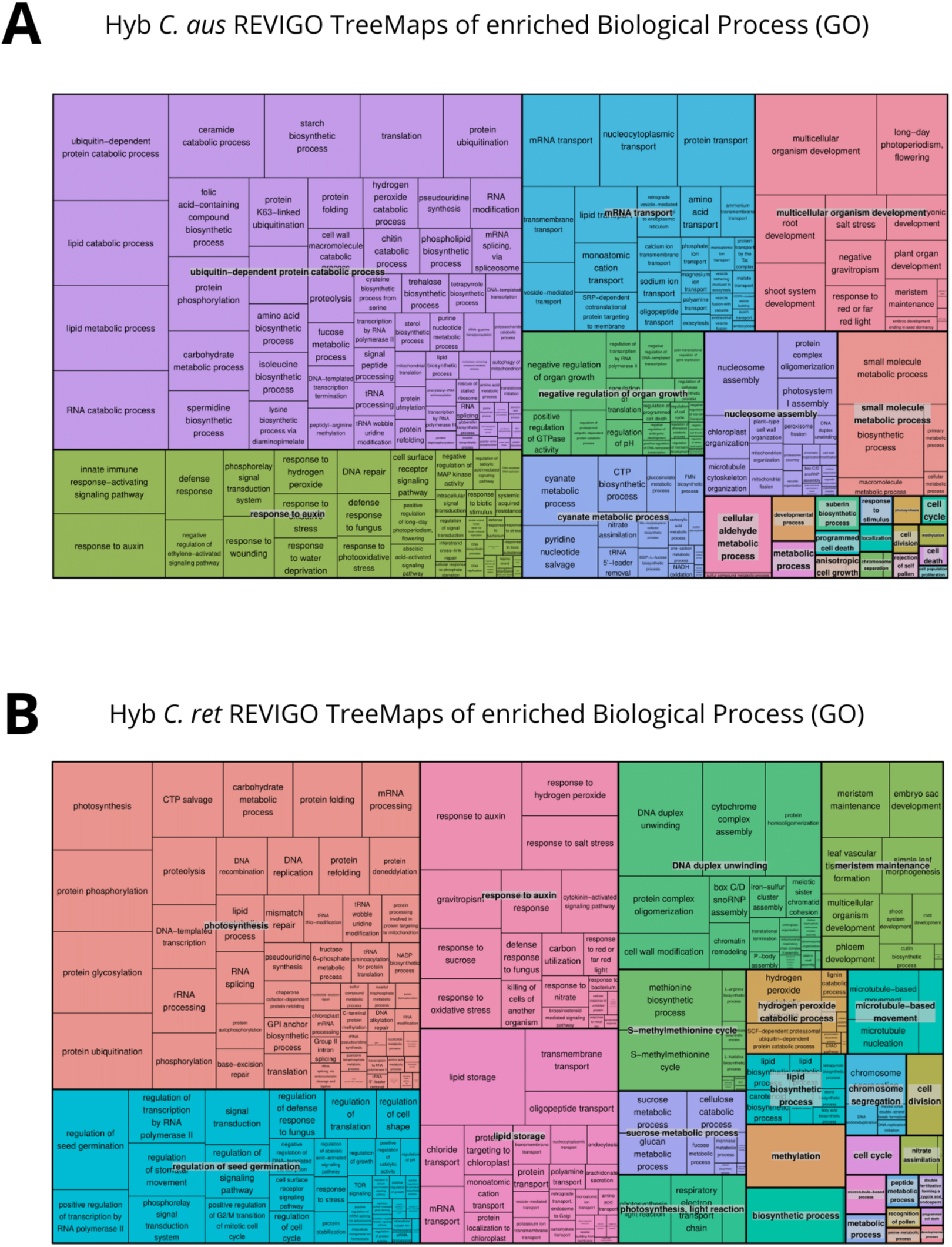
REVIGO TreeMaps of enriched Biological Process (GO) terms among differentially expressed genes (DEGs) in the hybrid. **(A)** GO terms enriched among genes upregulated from the C. australasica haplotype. Major categories include ubiquitin-dependent protein catabolic process, protein catabolic/metabolic process, translation, RNA modification and transport, and stress and defence responses. **(B)** GO terms enriched among genes upregulated from the C. reticulata haplotype. Enriched categories include photosynthesis, protein phosphorylation, RNA processing, DNA replication, NADP biosynthetic process, and other central metabolic and transcriptional functions, indicating a predominant maternal contribution to core cellular activities.

**Figure S4.**
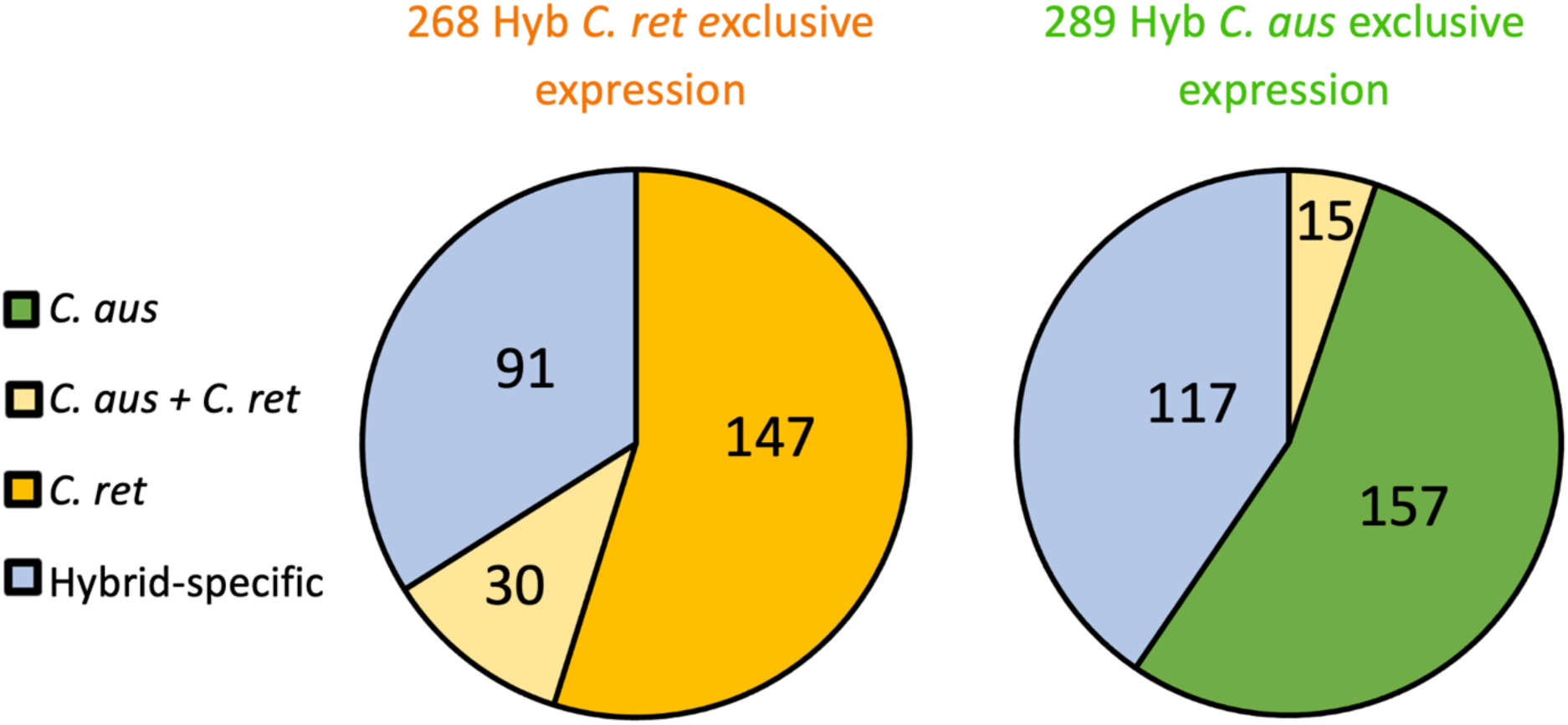
Exclusive allelic expression in the hybrid. Pie-chart diagram showing the origin of the allele exclusively expressed by the C. australasica haplotype (right) or the C. reticulata haplotype (left) in the hybrid. These genes are then separated in four categories: those that are only expressed in the hybrid and not in the parental lines (Hybrid-specific; blue), those that are also expressed in C. australasica (C. aus; green) or in C. reticulata (C. ret; orange) parental lines and those that are expressed in both parents (C. aus + C. ret; beige colour).

**Figure S5.**
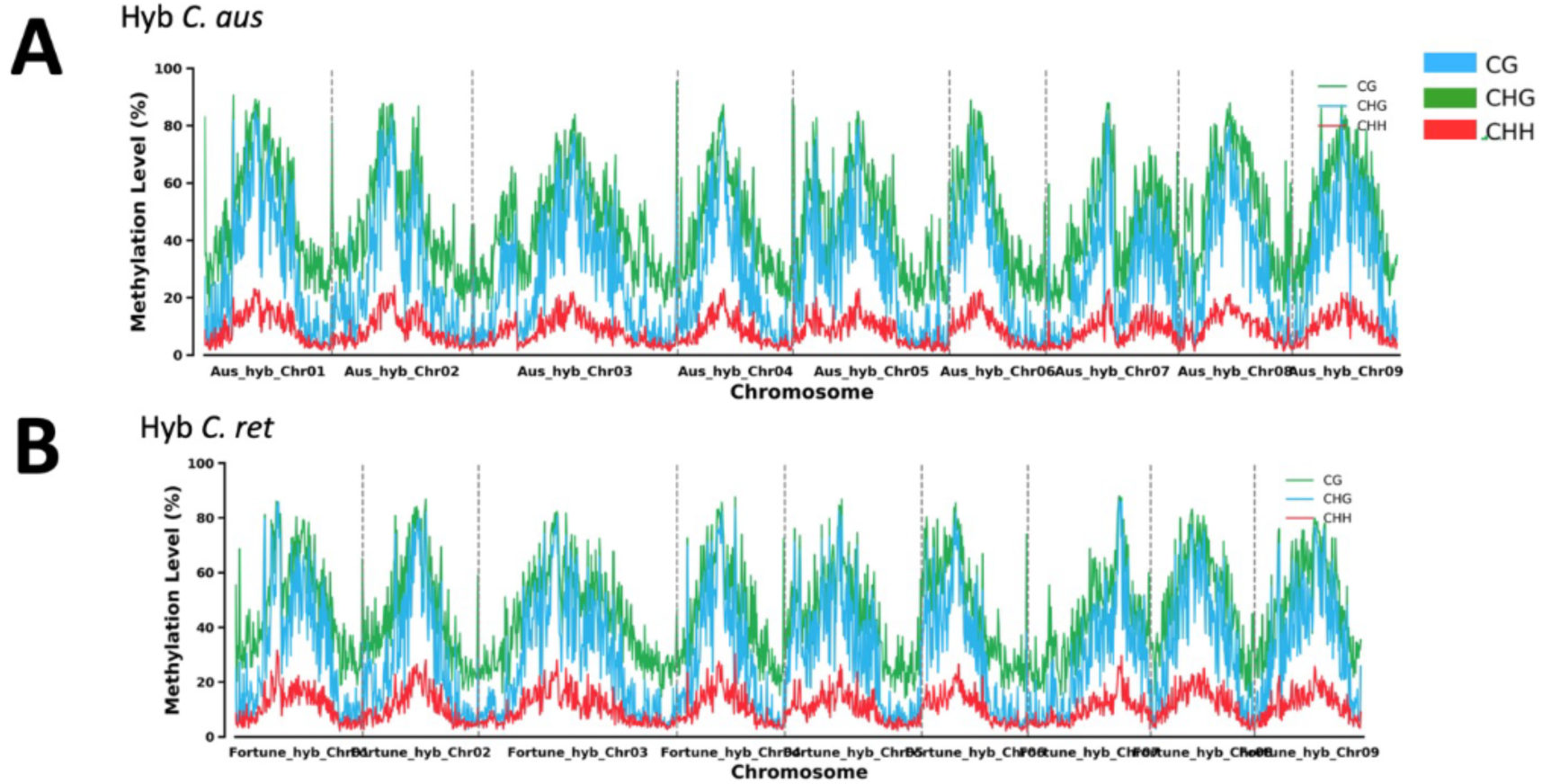
Genomic distribution of DNA methylation in the hybrid. The distribution of the three DNA methylation patterns (CG, CHG and CHH) is shown across all haplotype resolved hybrid C. aus (**A**) or hybrid C. ret. (**B**) chromosomes.

**Figure S6.**
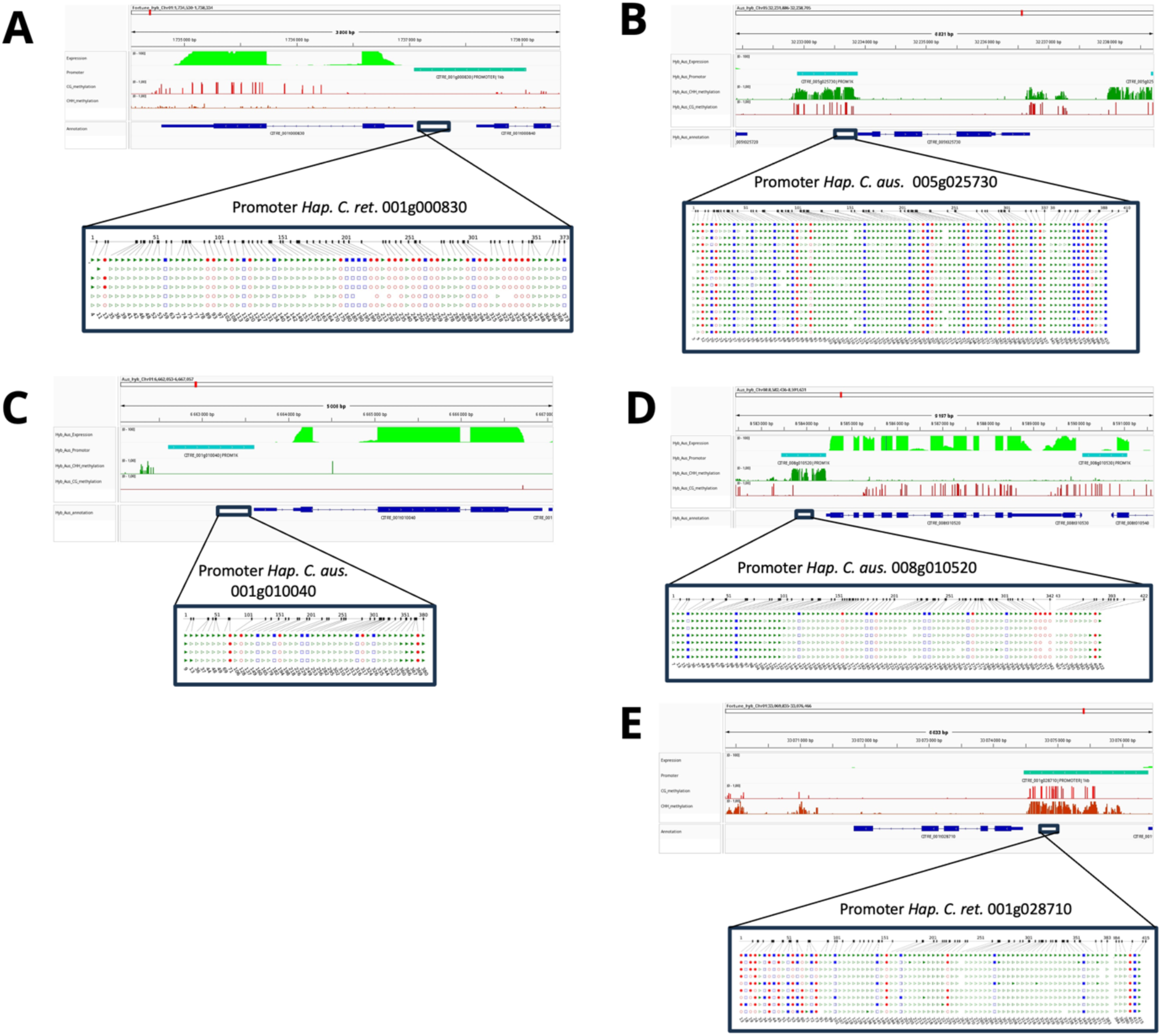
WGBS validation using BS-PCR and Sanger sequencing. To validate the methylation profiles obtained by WGBS, we designed primers at five different genomic location, performed bisulfite treatment and amplified selected promoter regions using specific primers. Browser screenshot displays the data obtained using WGBS and RNA-seq data, separating CHH and CG methylation. The black square corresponds to the genomic location amplified by PCR, cloned and sequenced using Sanger technic. Methylation analyses were then performed using CYMATE (www.cymate.org). As expected, unmethylated promoter regions in WGBS are also unmethylated when analysed by BS-PCR (**A**, **C**), while methylated promoter regions accumulate methylation in CHH context only (**D**), or at both CG and CHH context (**B**, **E**).

**Figure S7.**
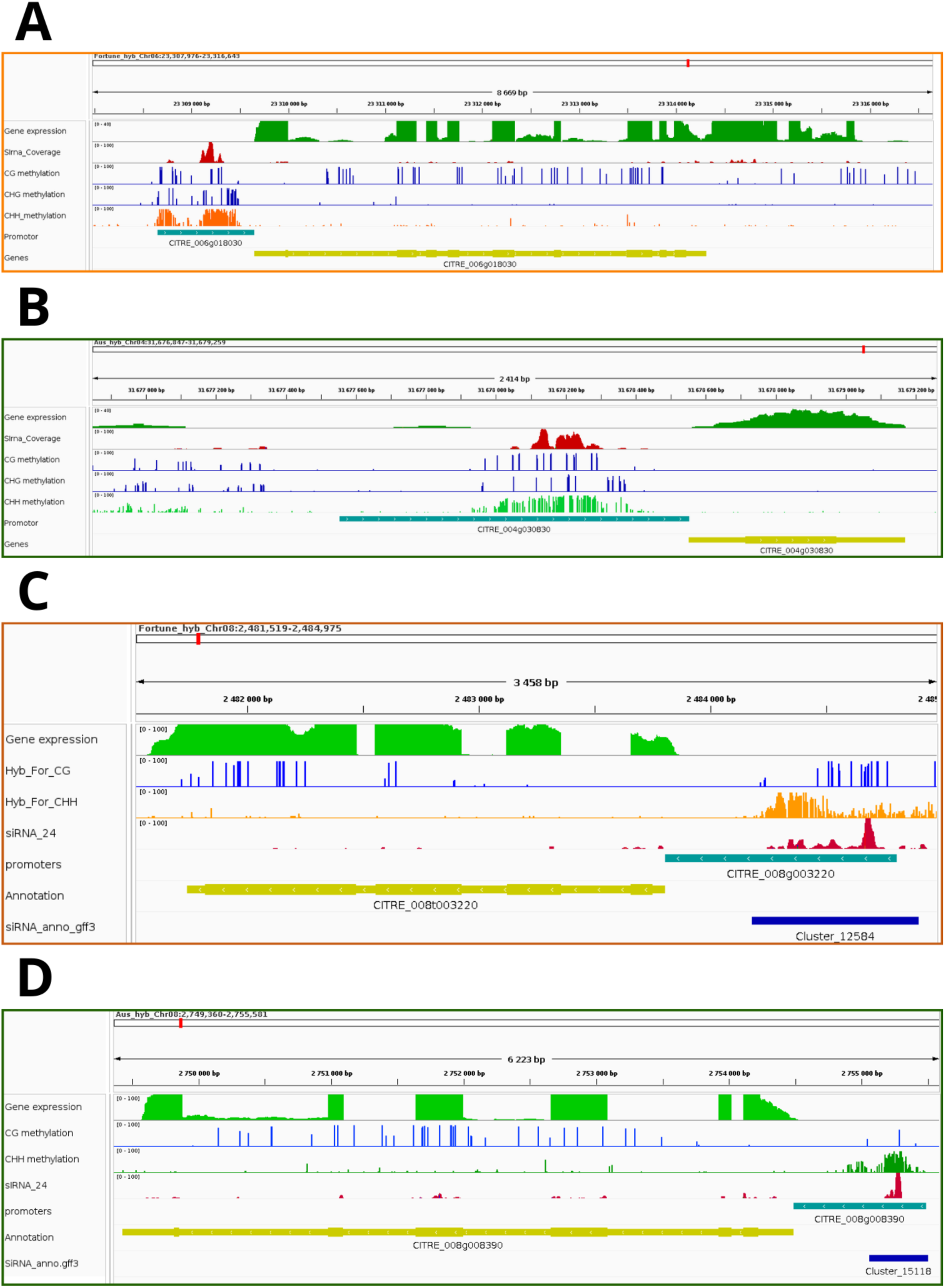
screenshot of IGV browser. IGV screenshot examples showing the distribution of small RNA clusters, mRNA accumulation, and CG or CHH methylation accumulation along 4 genomic loci in the hyb C. ret (**A, C**) or Hyb C. aus (**B, D**). In these 4 cases, the presence of siRNA clusters and mCHH in the promoter regions is not incompatible with gene expression.

**Figure S8.**
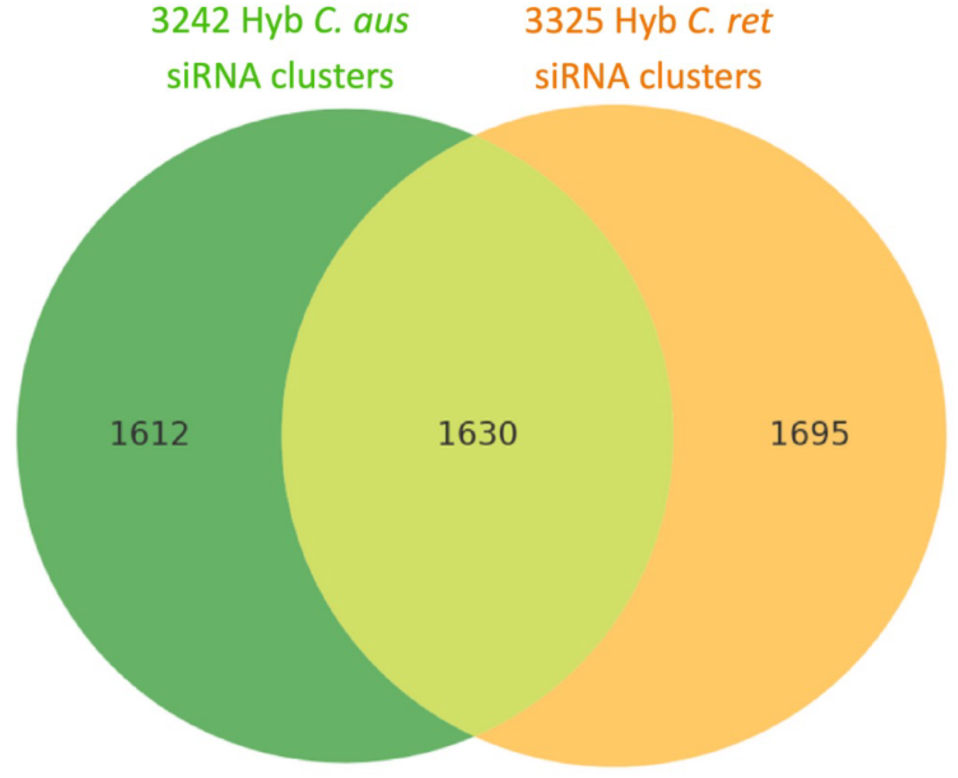
SiRNA clusters in the hybrid. Venn diagram showing the overlap of unique 24-nt siRNA sequences identified from clusters annotated independently in each haplotype (C. reticulata and C. australasica). About one third of 24-nt siRNA sequences (1630) are shared between the two haplotypes, while the remaining sequences are unique to one or the other: 1612 being specific to C. australasica haplotype (left, green), and 1695 being specific to the C. reticulata (right, orange) haplotype.

**Figure S9.**
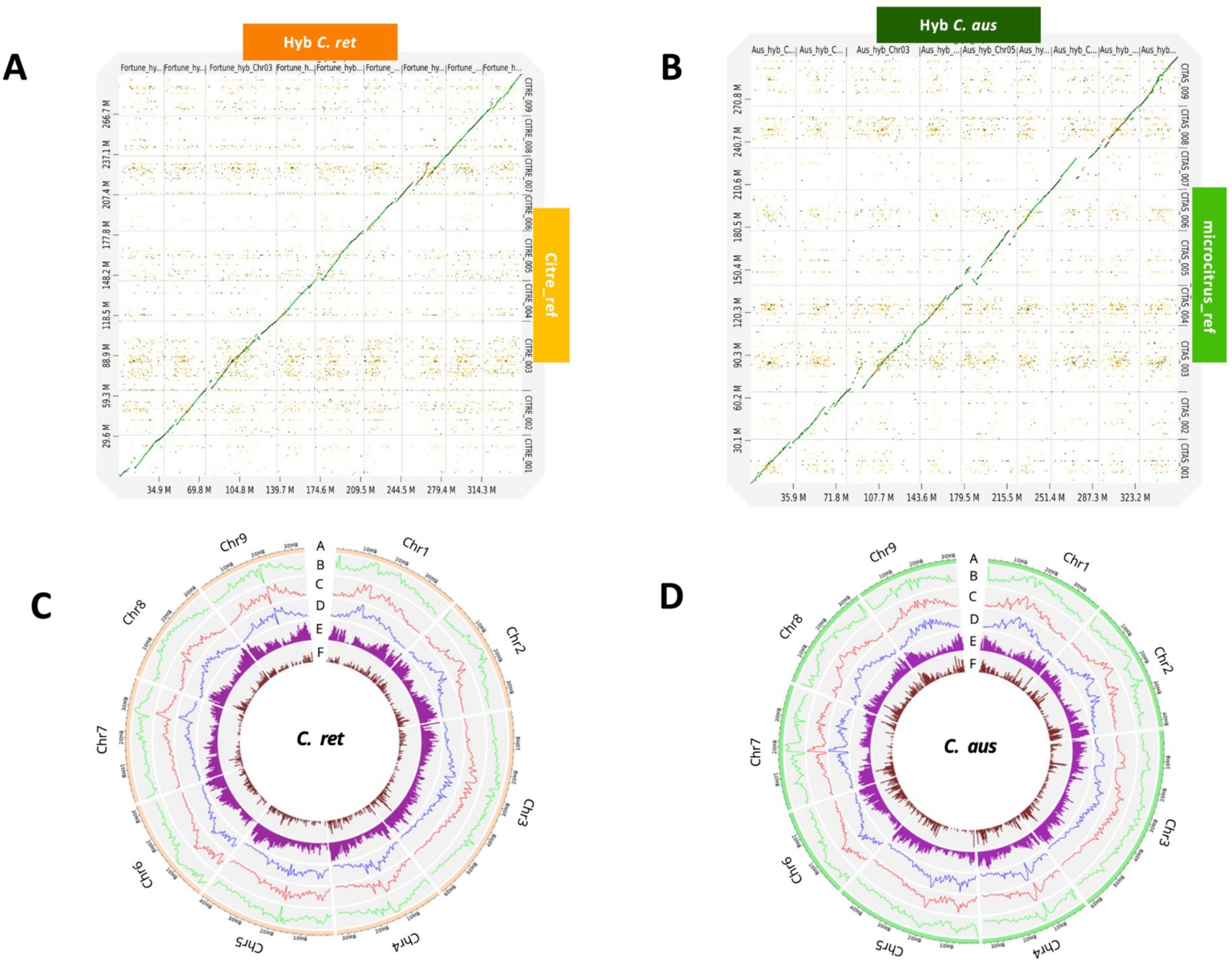
Genome-wide epigenomic patterns and overlap with expression in the parental lines. (**A**) Dot-plot alignment between the maternal haplotype of the hybrid C. ret and CITRE reference. **(B**) Dot-plot alignment between the paternal haplotype hybrid C. aus and the C. australasica reference genome. These plots illustrate overall synteny and structural rearrangements between each haplotype of the hybrid and the closest available reference genome from the corresponding parental line. Diagonal lines indicate conserved homologous regions, while off-diagonal signals highlight structural variations. (**C–D**) Circos plots representing genome-wide profiles of DNA methylation, gene density, and 24-nt siRNA clusters across the nine chromosomes of the two parental lines C. reticulata (**C**) and C. australasica (**D**).

**Figure S10.**
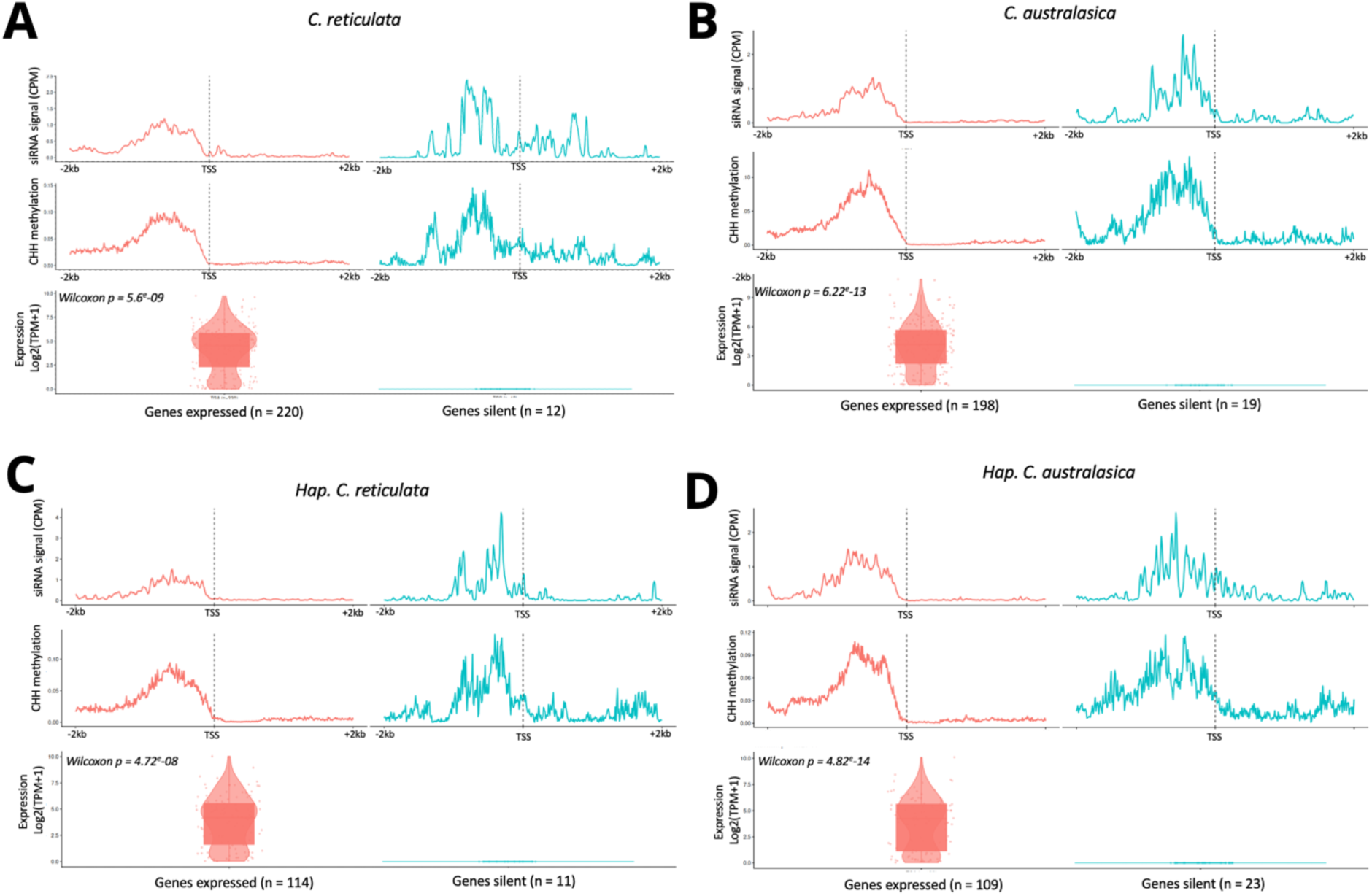
Specific analyses of siRNA–CHH methylation–gene expression modules in the C. ret, C. aus and in the hybrid. Metagene plots displaying 24-nt siRNA accumulation (top), CHH methylation (middle panel) and gene expression of siRNA–CHH methylation–gene modules with expressed genes (left panel) or silent genes (right panel) in C. reticulata (**A**), in C. australasica (**B**), or in the hybrid with an haplotype resolution (**C-D**). Metagene plots displays the accumulation from -2kb to +2kb of the Transcription start site (TSS).

**Figure S11.**
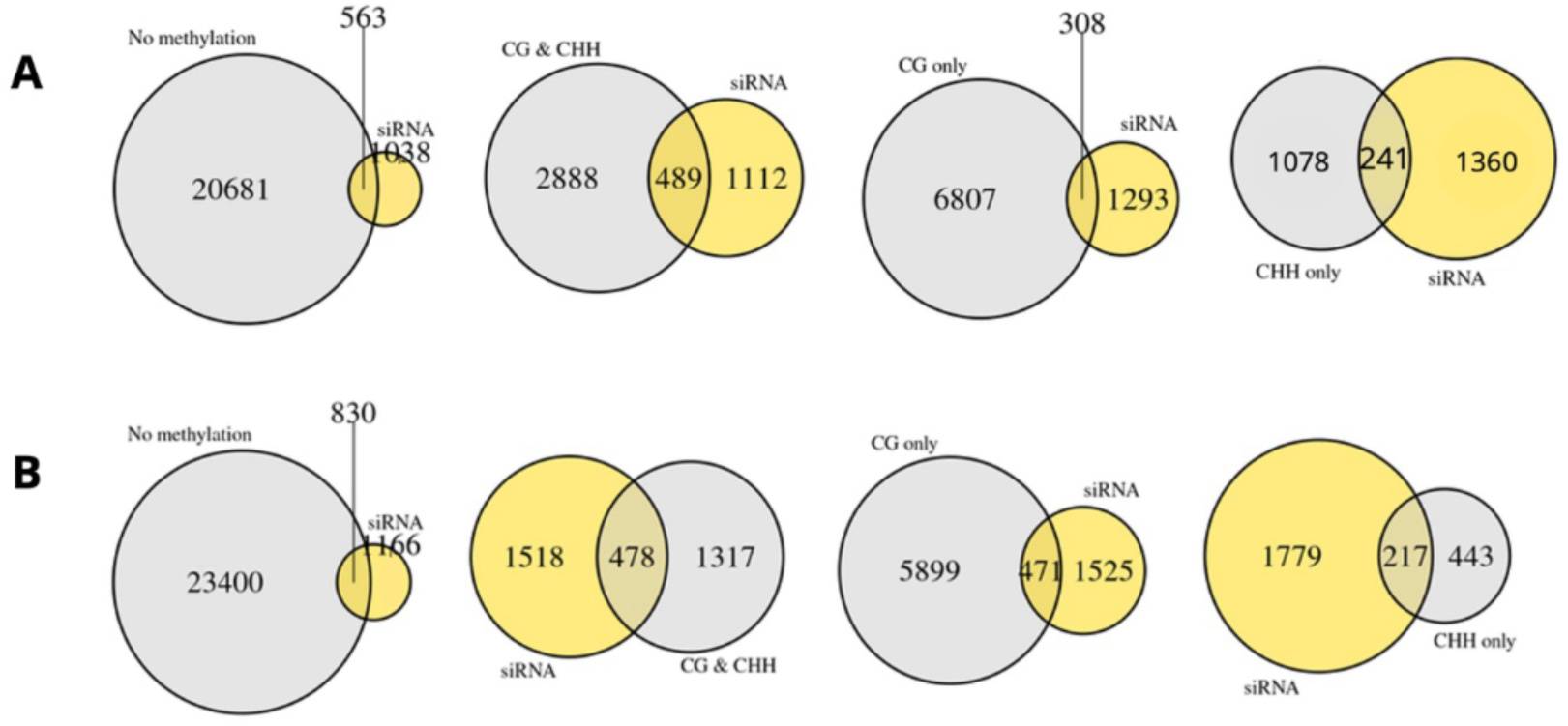
Venn diagrams showing the overlap between siRNA presence and methylation categories in parental genomes. Each plot displays the number of genes in a given methylation class (grey) and those with 24-nt siRNA clusters in their promoters (yellow) in C. reticulata (**A**) or in C. australasica (**B**). Overlaps represent genes with both features. Separate diagrams are shown for each methylation category.

**Figure S12.**
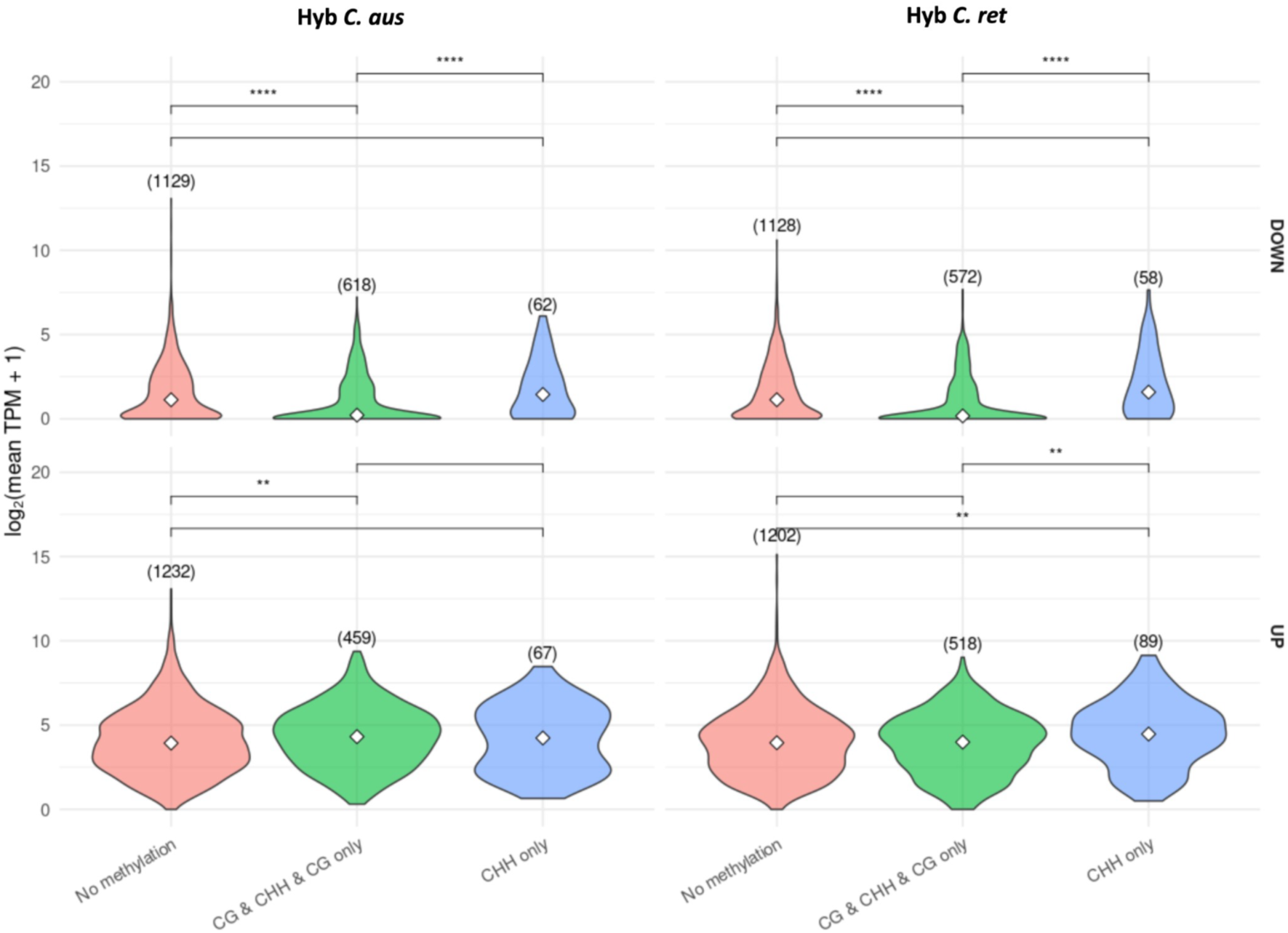
DNA methylation associating with the promoter of DEG in the hybrid. Violin plots displaying the distribution of DEG genes for each haplotype according to the type of methylation accumulating in their promoter regions. CHH > 20%; CG > 40%.

**Figure S13.**
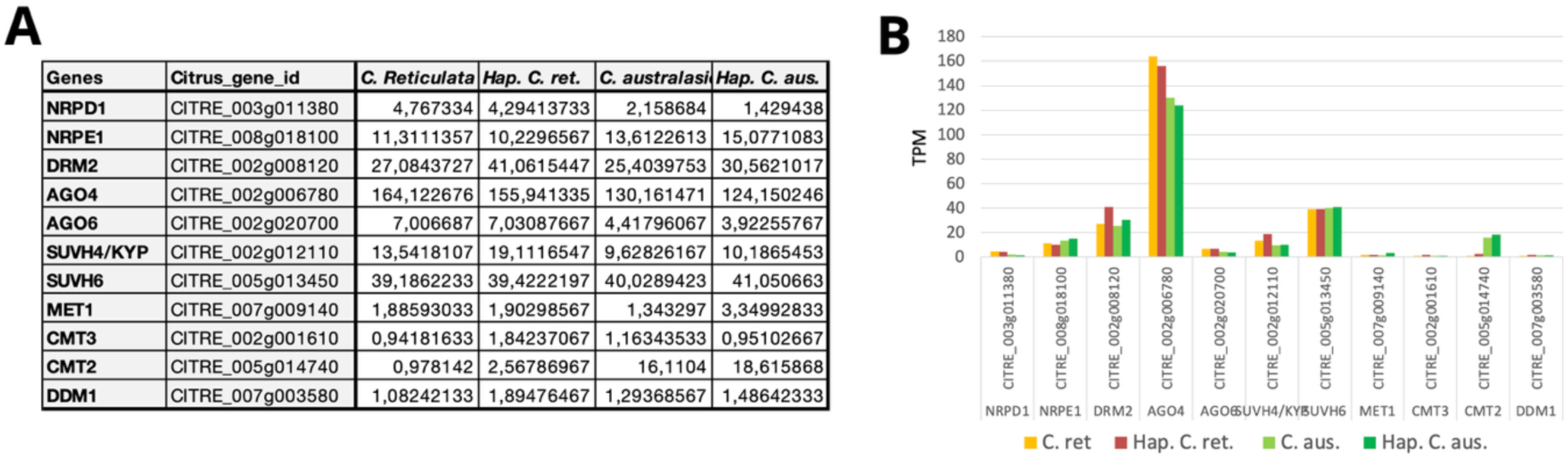
Relative expression of RdDM members in the hybrid and the parent. Analyses of the relative expression levels of several major homologs of the RdDM in the hybrid haplotypes and in the parental lines. Data were extracted from the RNA-seq and are presented as table (**A**) and in the graphic **(B**). TPM = Transcripts Per Kilobase Million

**Table S1.**
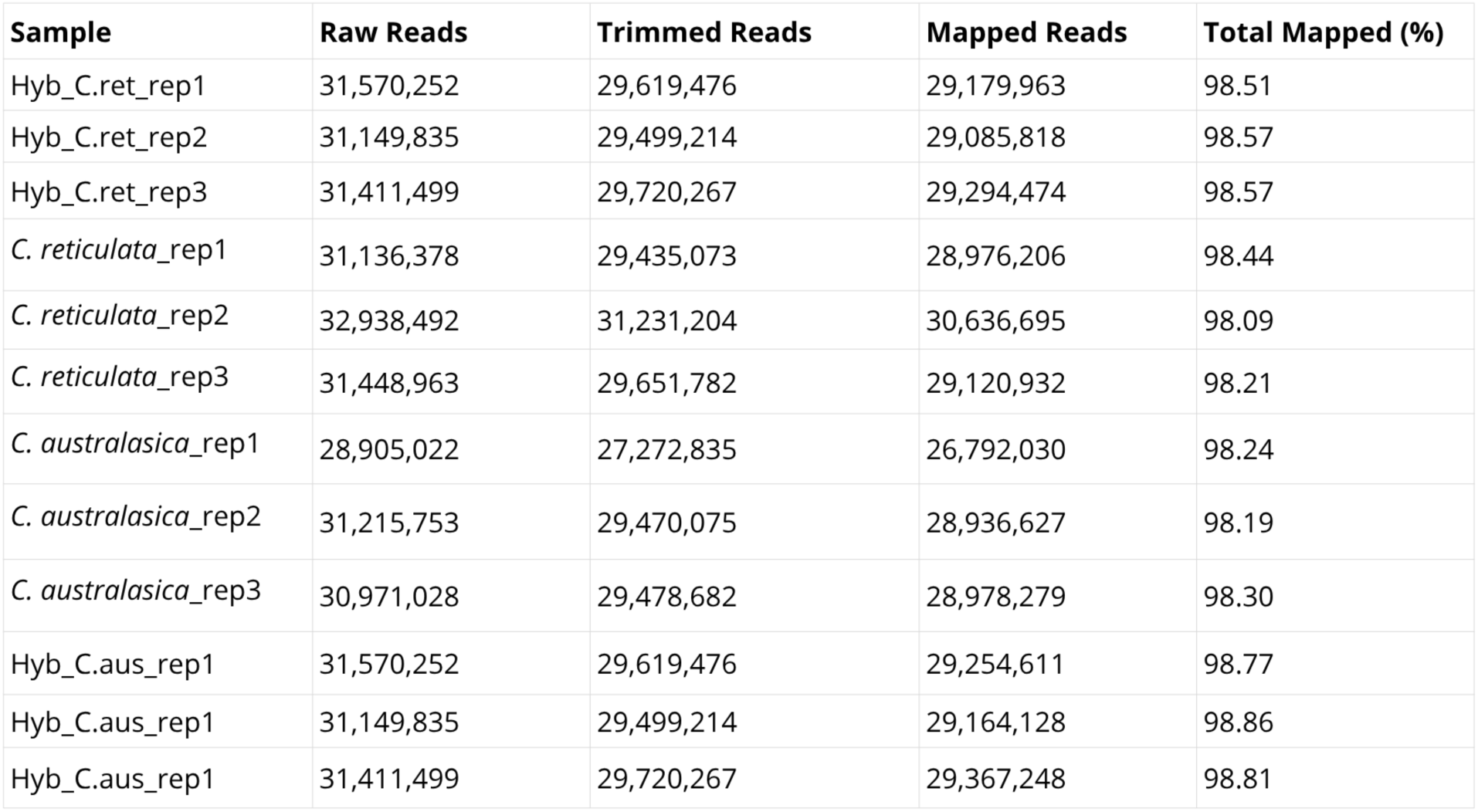
Summary of RNA-seq data processing and mapping. This table presents, for each replicate, the total number of raw reads, the number of reads retained after trimming, the number of mapped reads, and the total mapping rate (%). All samples exhibit high mapping rates (>98%), confirming the high quality of the RNA-seq libraries and the reliability of gene expression quantification in hybrid and parental genotypes.

| Sample | Raw Reads | Trimmed Reads | Mapped Reads | Total Mapped (%) |
| --- | --- | --- | --- | --- |
| Hyb_C.ret_rep1 | 31,570,252 | 29,619,476 | 29,179,963 | 98.51 |
| Hyb_C.ret_rep2 | 31,149,835 | 29,499,214 | 29,085,818 | 98.57 |
| Hyb_C.ret_rep3 | 31,411,499 | 29,720,267 | 29,294,474 | 98.57 |
| *C. reticulata*_rep1 | 31,136,378 | 29,435,073 | 28,976,206 | 98.44 |
| *C. reticulata*_rep2 | 32,938,492 | 31,231,204 | 30,636,695 | 98.09 |
| *C. reticulata*_rep3 | 31,448,963 | 29,651,782 | 29,120,932 | 98.21 |
| *C. australasica*_rep1 | 28,905,022 | 27,272,835 | 26,792,030 | 98.24 |
| *C. australasica*_rep2 | 31,215,753 | 29,470,075 | 28,936,627 | 98.19 |
| *C. australasica*_rep3 | 30,971,028 | 29,478,682 | 28,978,279 | 98.30 |
| Hyb_C.aus_rep1 | 31,570,252 | 29,619,476 | 29,254,611 | 98.77 |
| Hyb_C.aus_rep1 | 31,149,835 | 29,499,214 | 29,164,128 | 98.86 |
| Hyb_C.aus_rep1 | 31,411,499 | 29,720,267 | 29,367,248 | 98.81 |

**Table S2.** Mapping statistics of WGBS data for hybrid and parental samples. For each biological replicate, the table reports the number of paired-end reads obtained, the number of uniquely aligned pairs, the mapping efficiency (%), the duplication rate (%), the number of deduplicated reads, the estimated genome size used for mapping (in megabases), and the average sequencing depth (X). The samples include the hybrid mapped to both parental haplotypes (Hyb_C.ret and Hyb_C.aus), as well as the two parental species (C. reticulata and C. australasica).

| Samples | Sequence pairs | Number of paired-end alignments with a unique best hit | Mapping Efficiency (%) | Duplication Rate (%) | Deduplicated Leftover Sequences | Genome Size (Mb) | Average Depth (X) |
| --- | --- | --- | --- | --- | --- | --- | --- |
| Hyb_C.aus_rep1 | 59471997 | 45299369 | 76.2 | 21.17 | 35707603 | 359 | 25.56 |
| Hyb_C.aus_rep2 | 56997431 | 43090193 | 75.6 | 23.2 | 33092958 | 359 | 24.5 |
| Hyb_C.aus_rep3 | 52490561 | 40235003 | 76.7 | 22.88 | 31029064 | 359 | 22.56 |
| Hyb_C.ret_rep1 | 59471997 | 44722620 | 75.2 | 21.75 | 34997498 | 349 | 25.56 |
| Hyb_C.ret_rep2 | 56997431 | 42539674 | 74.6 | 23.69 | 32461335 | 349 | 24.5 |
| Hyb_C.ret_rep3 | 52490561 | 39753625 | 75.7 | 23.55 | 30389785 | 349 | 22.56 |
| *C. reticulata*_rep1 | 54120503 | 43591838 | 80.5 | 23.67 | 33274917 | 349 | 23.96 |
| *C. reticulata*_rep2 | 53416001 | 41156597 | 77.0 | 22.69 | 31818808 | 349 | 22.82 |
| *C. reticulata*_rep3 | 50019636 | 38564496 | 77.1 | 18.48 | 31436564 | 349 | 22.67 |
| *C. australasica*_rep1 | 53564928 | 41617819 | 77.7 | 26.86 | 30441058 | 359 | 21.67 |
| *C. australasica*_rep2 | 54127489 | 41896425 | 77.4 | 26.90 | 30627038 | 359 | 21.81 |
| *C. australasica*_rep3 | 53798201 | 41649738 | 77.4 | 25.70 | 30946374 | 359 | 21.97 |

**Table S3.**
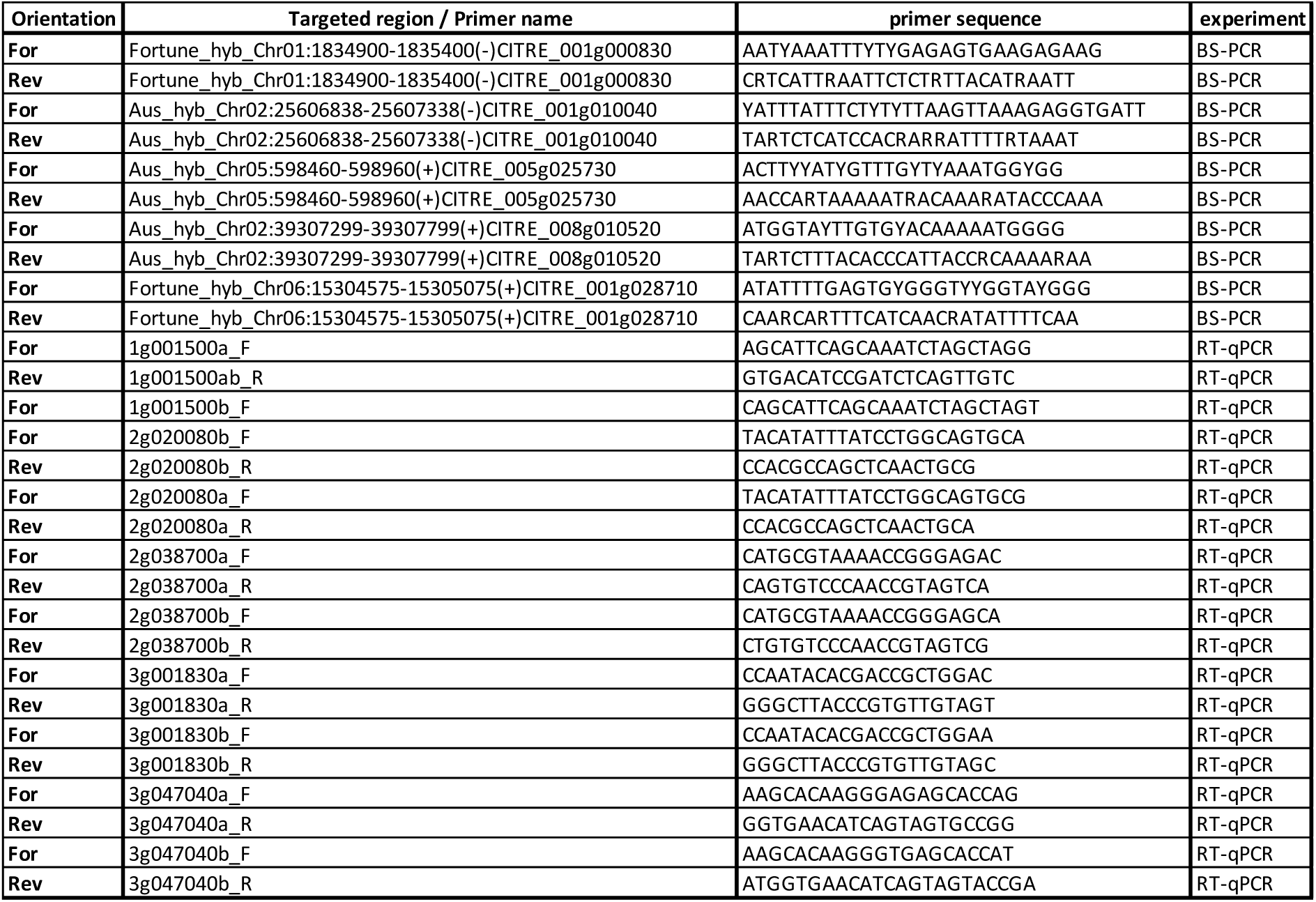
List and sequences of the primer used in this study.

