## Supplemental Material and method for "Investigation of allele-specific expression in citrus hybrids reveals the association between siRNA-mediated de novo methylation and high expression in citrus genomes"

### Supporting information on MATERIALS AND METHODS

#### Nucleic Acid Extraction

Leaf tissues of *C. reticulata*, *C. australasica* of the F1 hybrid were ground in liquid nitrogen using a pre-chilled mortar and pestle.

***DNA Extraction for Nanopore Sequencing***: High-molecular-weight genomic DNA was extracted from 0.7 g of ground leaf tissue using the "High Molecular Weight Genomic DNA for Long-Read NGS" protocol recommended by Oxford Nanopore Technologies.

***DNA Extraction for WGBS***: Genomic DNA was extracted using the DNeasy Plant Mini Kit (Qiagen) according to the manufacturer's instructions.

***RNA Extraction***: Total RNA was extracted from 70–80 mg of ground leaf tissue using the kit RNeasy Plant Mini Kit (Qiagen) following the manufacturer's instructions.

Nucleic acid concentrations and purity were assessed using a Nanodrop 2000 (Thermo Scientific) and Qubit Fluorometer (Thermo Fisher Scientific).

#### Sequencing

***Tell-Seq Sequencing***: Genomic DNA from the two parental genotypes *Citrus reticulata* and C*itrus australasica* were sequenced using Tell-Seq technology (Universal Sequencing Technology), Tell-seq libraries were prepared for these two accessions according manufacturer's instructions and pair-end sequenced using Illumina Novaseq platform at Montpellier Genomix with an average coverage of 30X per accession. In this protocol, long DNA fragments are partitioned and barcoded within microreactors prior to Illumina short-read sequencing (2 × 150 bp), enabling the reconstruction of synthetic long reads from paired-end data. These data were used to extract haplotype-specific k-mers for the trio-binning step, allowing assignment of Nanopore reads from the hybrid to their maternal or paternal origin.

***Short-Read Sequencing***: RNA-seq, WGBS, and small RNA libraries were prepared and sequenced by Novogene. RNA-seq and WGBS libraries were sequenced using Illumina 2 × 150 bp paired-end reads. Small RNA libraries were sequenced using Illumina 50 bp single reads.

**Scaffolding and reference-based ordering:** Contigs were ordered with RagTag [1] using *C. reticulata* (*CITRE* [2]) and *C. australasica (CITAS,* [3]*)* genome assemblies as references.

**Assembly quality assessment:** The quality of the scaffolded haplotypes was assessed with QUAST v5.0 [4] and BUSCO v5 [5] using the eudicotyledons_odb10 orthologous gene set and default parameters.

**Genome anchoring on Citrus consensus mapping:** To assess the collinearity between haplotype assemblies and the Citrus consensus genetic map, markers were anchored using the locOnRef tool from the Scaffhunter toolbox [6]. The reference map includes 10,756 loci, comprising 7,915 gene-based markers and 2,841 non-genic SNPs [7]. Alignments were performed with blastn [8] (identity ≥ 95%, length ≥ 100 bp), retaining only unique best hits per marker. Anchoring quality and genome collinearity were visualized using Circos [9]**.**

**Visualization of genome alignments:** Dot plots comparing the scaffolded haplotypes and the reference genomes were generated using D-GENIES [10].

**RNA-seq Analysis**

Raw RNA-seq reads were quality-checked using FastQC and trimmed with Trimmomatic. cleaned reads were aligned to each haplotype genome using HISAT2 v2.2.1 [11] (Table S1). Alignments were processed using SAMtools v1.10 [12]. Gene expression quantification was performed using featureCounts (Subread package) [13]. Differential expression analysis was conducted with DESeq2 [14] in R, considering genes with adjusted p-values ≤ 0.01 and |log2FoldChange| ≥ 1.5 as differentially expressed. Gene Ontology enrichment analysis was performed using topGO [15] , and significant terms were visualized with REVIGO [16].

**Whole-Genome Bisulfite Sequencing (WGBS) Analysis**

WGBS reads were aligned to haplotype-resolved genomes using Bismark v0.24.1 [17] (Table S2). Duplicate reads were removed using deduplicate_bismark. Methylation levels at CG, CHG, and CHH contexts were extracted using coverage2cytosine and bismark_methylation_extractor. Methylation profiles around genes (±2 kb) were generated using ViewBS [18]. Genome-wide methylation patterns were visualized using MethGo [19] and shinyCircos-V2.0 [20], highlighting regional and haplotype-specific differences.

**Small RNA Analysis**

Cleaned small RNA reads (18–30 nt) provided by the sequencing platform were analysed using ShortStack v3.8.5 [21] for alignment and cluster identification with default options. Clusters were categorized based on dominant read length, focusing on 24-nt siRNA clusters. Briefly, clusters are formed based on local sRNA coverage, and any genomic position with a coverage above the minimum threshold (1 RPM) is considered as a potential peak. Peaks located within 200 nt of each other are then merged into a single siRNA locus. Examples of siRNA clusters can be visualized *via* IGV (Integrative Genomics Viewer) [22], screenshots (Suppl. Figures). Clusters intersecting with promoter regions (1 kb upstream of transcription start sites) were identified using GFF3 outputs from ShortStack. Comparative analyses identified haplotype-specific and shared clusters. These clusters were integrated with differential expression and methylation data to explore potential regulatory interactions.

**Bisulfite-PCR (BS-PCR)**

Bisulfite treatment was performed using the EZ DNA Methylation (Zymo Reseach, Ref D5001) following manufacturer's instructions and as previously described [23] and PCR primers described in the (Table S3). Analyses were then performed using Cymate (https://cymate.org/) [24].

**RT-qPCR**

RT-qPCR were perfromed as previously described in (Jarry et al., 2023) [25]. Primers used are described in the Table S3.

**Statistical Analysis**

All tests were conducted in R (v4.5.0). Gene TPMs were $\log_2$-transformed ($\log_2$[mean TPM + 1]).

***Methylation vs. expression***: Promoters were binned into four categories (No methylation, CHH-only, CG-only, CG & CHH) using thresholds of 20 % CHH and 40 % CG. Expression differences among categories were tested by Kruskal–Wallis; significant global results ($p < 0.05$) were followed by Dunn's post-hoc tests with Benjamini–Hochberg correction.

***siRNA presence vs. expression:*** Promoters were labelled "with" or "without" siRNA 24-nt clusters. $\log_2$(TPM) distributions were compared by Wilcoxon rank-sum tests (two-sided), with p-values adjusted (BH) across origins.

***Spearman's rank correlation:*** Two approaches were used: (1) we averaged expression values across replicates and compared the mean expression for each gene, and (2) we performed the correlation analysis independently for each replicate. Both approaches yielded consistent results.
